# Maresin 1 Repletion Improves Muscle Regeneration After Volumetric Muscle Loss

**DOI:** 10.1101/2022.11.19.517113

**Authors:** Jesus A. Castor-Macias, Jacqueline A. Larouche, Emily C. Wallace, Bonnie D. Spence, Alec Eames, Benjamin A. Yang, Carol Davis, Susan V. Brooks, Krishna Rao Maddipati, James F. Markworth, Carlos A. Aguilar

## Abstract

The acute traumatic or surgical loss of skeletal muscle, known as volumetric muscle loss (VML), is a devastating type of injury that results in exacerbated and persistent inflammation followed by fibrosis. The mechanisms that mediate the magnitude and duration of the inflammatory response and ensuing fibrosis after VML remain understudied and as such, the development of regenerative therapies has been limited. To address this need, we profiled how lipid mediators, which are potent regulators of the immune response after injury, varied with VML injuries that heal or result in fibrosis. We observed that non-healing VML injuries displayed increased pro-inflammatory eicosanoids and a lack of pro-resolving lipid mediators. Treatment of VML with a pro-resolving lipid mediator synthesized from docosahexaenoic acid, called Maresin 1, ameliorated fibrosis through reduction of neutrophils and macrophages and improved myogenesis, leading to enhanced recovery of muscle strength. These results expand our knowledge of the dysregulated immune response that develops after VML and identify a novel immuno-regenerative therapeutic modality in Maresin 1.

## INTRODUCTION

The acute loss of a large volume of skeletal muscle, or volumetric muscle loss (VML), is a severe type of injury that results extensive fibrosis and attenuation of muscle function(1) and disability(2). Despite attempts to use regenerative medicine and tissue engineering approaches(3), VML injuries typically result in sustained inflammation, supplantation of muscle tissue with excessive extracellular matrix(4), and muscle fibrosis(5, 6). The cellular and molecular drivers that recruit and program the fibrotic response(7) after VML remain understudied. As such, regenerative therapies to restore muscle function after VML have found limited success.

A coordinated immune response to injury is critical for muscle repair, and begins with a pro-inflammatory state that is characterized by leukocyte infiltration followed by an anti-inflammatory phase and regenerative response of resident stem and progenitor cells(8, 9). An impairment in the transition from a pro-inflammatory immune phenotype towards a pro-repair state leads to secondary damage and inefficient regeneration(8). VML has been shown to result in exacerbated infiltration and/or impaired clearance of neutrophils(10), but the role of inflammatory macrophages(11, 12) and changes in their polarization remain understudied. Macrophages are critical regulators of tissue repair(13), and delays or interruptions of their transition away from a pro-inflammatory state(14, 15), lead to adoption of a fibrotic, TGFβ1-secreting phenotype(12). There is a *void* in our understanding of macrophage dysfunction that develops after VML and the causative extracellular signaling changes that confer this pathological response.

Recently, a new class of bioactive signaling factors derived from omega-3 or essential fatty acids called pro-resolving lipid mediators(16) have been discovered to regulate the magnitude and duration of the inflammatory response(17). These pro-resolving lipid mediators have been shown to restrain the infiltration of neutrophils(18), augment macrophage polarization(19) and phagocytosis(20), and attenuate pro-inflammatory signaling molecules, such as eicosanoids(21). The balance that develops after VML between classical inflammatory eicosanoids derived from arachidonic acid (e.g. prostaglandins and leukotrienes(22, 23) and pro-resolving lipid mediators such as resolvins, protectins, and maresins has not been evaluated. Moreover, how many of these lipid mediators contribute to recruitment and conditioning of immune cell subtypes after VML and concomitant fibrosis requires further understanding.

Herein, we contrasted VML injuries that heal and restore muscle function with those that result in fibrosis and loss of muscle function. We used metabolipidomics analysis over a time course to assess changes in the composition of bioactive signaling mediators for VML injuries that do not heal compared to those that result in fibrosis. For degenerative VML injuries, increased pro-inflammatory eicosanoids were detected when compared to VML injuries that heal and no detectable change in pro-resolving mediators. Exogenous administration of a docosahexaenoic acid-derived pro-resolving lipid mediator called Maresin 1(20, 24) after degenerative VML injury was observed to impact resolution trajectory by simultaneously attenuating macrophage and neutrophil infiltration, reducing fibrosis, and promoting muscle regeneration via enhancing muscle stem cell proliferation. These findings suggest pro-resolving lipid mediators can encourage healing of severe muscle trauma and alter the signaling environment to support muscle stem cell-based regeneration.

## RESULTS

### Comparative Analysis of Volumetric Muscle Loss Injuries of Varied Sizes Reveals Variations in Fibrosis and Function

To establish a framework for understanding how variations in the inflammatory response drive fibrotic scarring and muscle degeneration after VML injury, we administered bi-lateral VML injuries to the tibialis anterior (TA) muscles of adult C57BL6/J mice by delivering full thickness 1mm or 2mm punch biopsies(25) (Figure 1A). We extracted muscles at 7- and 14-days post injury (dpi) and observed increased collagen deposition in 2mm defects when compared to 1mm defects by Picrosirius red staining (Figures 1B-D, n=3-4 mice per condition, paired). To determine if the increases in fibrosis with larger VML defects engendered reductions in maximal tetanic force, we compared 1mm and 2mm defects with uninjured muscle at 28 dpi. We found reductions in force output for 2mm defects when compared to 1mm and uninjured muscle (Figures 1E-F, Supp. Fig. 1A-C, n=6-8 mice per group, unpaired two-way ANOVA), which is consistent with previous studies(10, 25). Summing these results shows that 2mm punch biopsy defects to murine TA muscles (degenerative VML injuries) produce fibrotic supplantation and reductions in muscle function, while 1mm punch biopsy defects (regenerative VML injuries) result in less fibrosis and functionally recover to the same level as uninjured tissues by 28 dpi.

**Figure 1.**
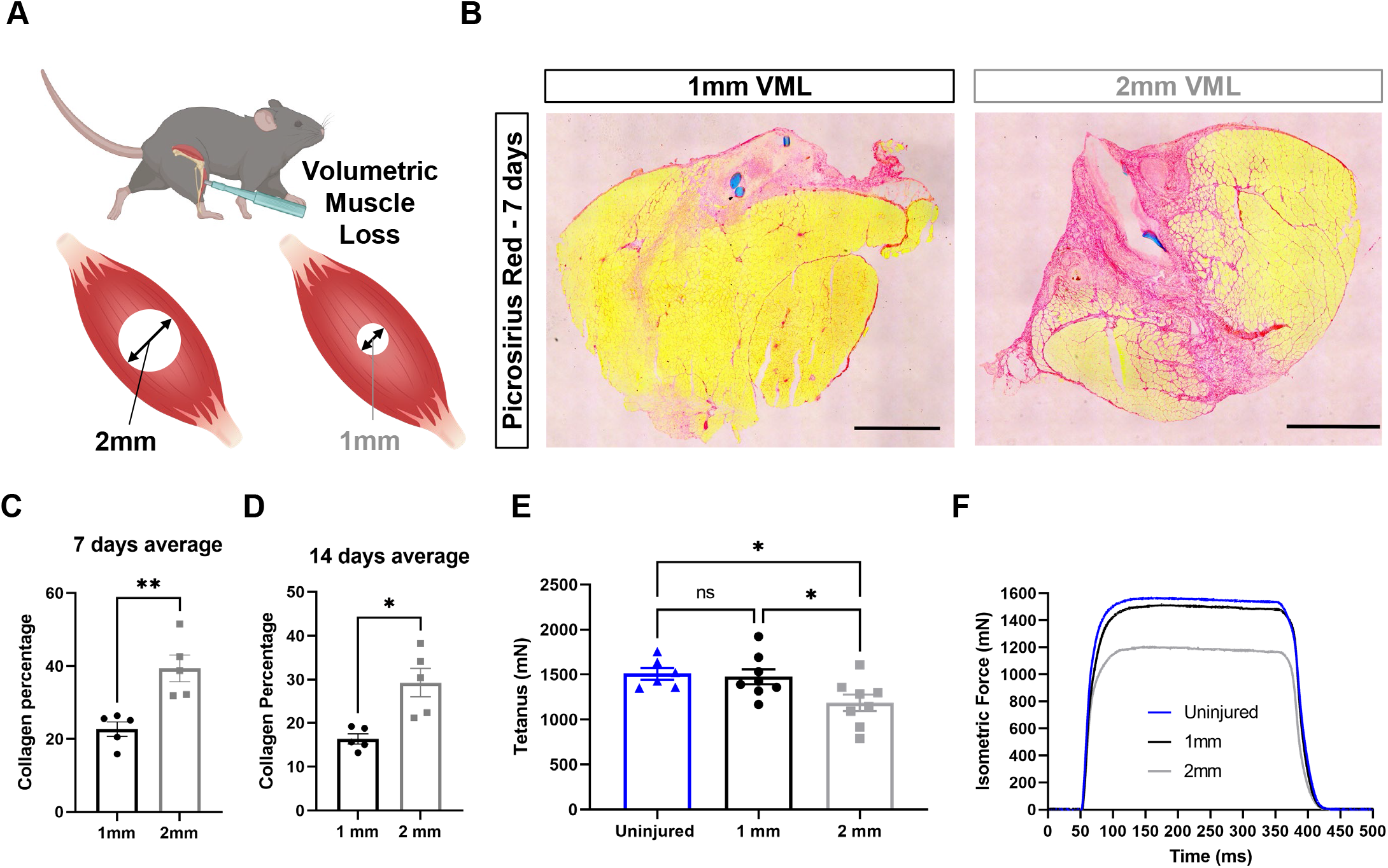
In vivo comparison of degenerative and regenerative volumetric muscle loss injuries. **A**) Schematic of experiment, whereby young (3 months) mice were administered 1mm or 2mm biopsy punches to their tibialis anterior muscle to create a volumetric muscle loss (VML) injury. **B)** Representative cross-sections stained with Picrosirius Red isolated from healing (1mm defect) and non-healing (2mm defect) 7 days after VML muscle injuries (n=4-5/group). Scale = 200μm. **C)-D)** Quantitation of images from B) show reductions in Picrosirius Red for 1mm defects compared to 2mm defects at 7 days (C) and 14 days (D). Graphs show mean ± SEM. **E)** Average tetanic force from uninjured (blue) tibialis anterior muscle at 28 days following 1mm (black) or 2mm (gray) VML injuries. Bars show mean ± SEM and *p<0.05 between injury types by two-way ANOVA and post-hoc. n=6-8 / group. **F)** Representative force curves of uninjured tibialis anterior muscle (blue) at 28 days following 1mm (black) or 2mm (gray) injuries. For C) and D), unpaired t-test with Welch’s correction. * p< 0.05 and ** p< .01.

### Metabolipidomic Profiling After Volumetric Muscle Injuries Shows Imbalances in Pro- and Anti-Inflammatory Lipid Mediators

A dysregulated immune response(7, 10) has been shown to be responsible for the fibrotic scarring induced from degenerative VML injury. To glean the factors that recruit and condition myeloid derived cells to promote excessive tissue fibrosis, we administered regenerative (1 mm) vs degenerative (2mm) VML injuries to TA muscles as above and performed liquid chromatography coupled to tandem mass spectrometry (LC-MS/MS) based metabolipidomics profiling at 0, 3, 7, and 14 dpi (Figure 2A). We profiled a total of 143 lipid mediator species across the time course of recovery from VML. In total, 80 lipid mediators were reliably detected in muscle tissue homogenates (signal to noise ratio >3 and peak quality >0.2 in at least 50% of samples).

**Figure 2.**
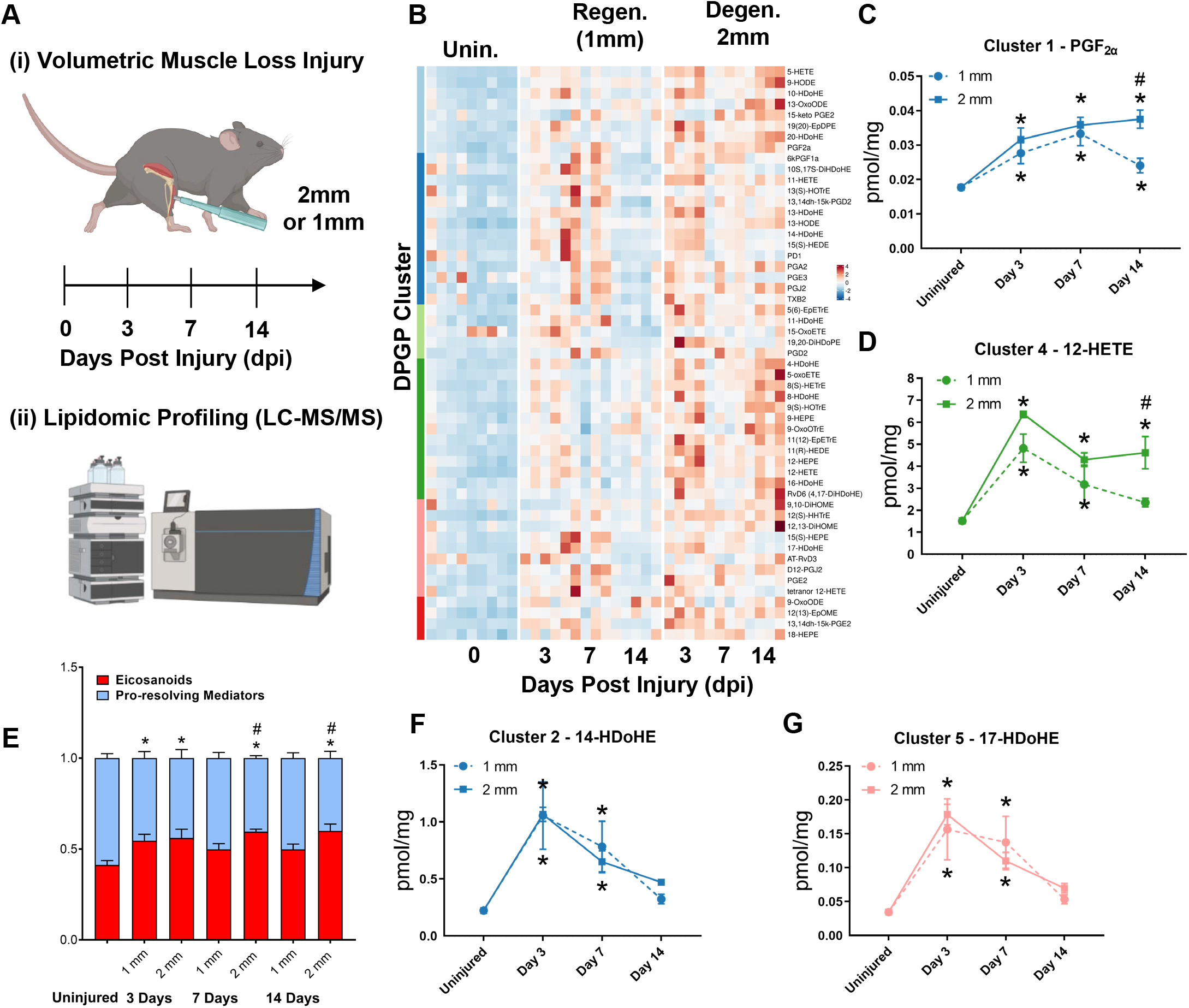
Lipidomic profiling after volumetric muscle loss injuries reveals stronger and sustained lipid mediator response in degenerative injuries. **A**) Mice were administered a bilateral defect to TAs (left leg received a 1mm defect and right leg received a 2 mm defect). Tissues were harvested at 0, 3, 7 and 14 days post injury and subjected to LC/MS-MS analysis. n=5 mice per injury type and timepoint. **B)** Row-scaled heatmap of 53 differentially expressed analytes across injuries and timepoints. Analytes are grouped by Dirichlet Process Gaussian Process (DPGP) clusters. **C-D, F-G)** Changes of specific lipid metabolites in different clusters per injury type and timepoint expressed in pmol/mg. Two-way paired ANOVA was used to estimate statistical differences between injury and timepoints. Graphs show mean ± SEM, * < 0.05 between timepoints of matched injury timepoint, ** < .01 between different injury types, and # < 0.05 between different injury types at a certain timepoint. **CD)** Prostaglandin F2 alpha and 12-HETE are both eicosanoids related to pro-inflammatory effects. **E)** Eicosanoids (TXB2, PGD2, PGE2, PGF2α, 6kPGF1α, and 5-, 12-, 15-HETEs) vs pro-resolving mediators pathway markers (5-HEPE, 4-HDoHE, 7-HDoHE, 12-HEPE, 14-HDoHE, RvD6, PD1, 10S,17S-DiHDoHE, Maresin 1, Mar1-n3DPA, LXA4) graphed for 2 mm and 1 mm volumetric muscle loss (VML) defects at day 0,3,7,14. Both analyte groups were normalized to ratios adding up to 1. Two-way paired ANOVA statistical tests were used to compare values between injury and timepoints. * < 0.05 between timepoints of matched injury timepoint, and # < 0.05 between different injury types at a certain timepoint **F)** 14-HDoHE represents a pathway marker for maresins. **G)** 17-HDoHE is a pathway marker for D-resolvins/protectins. Both Maresins and Resolvins are related to anti-inflammatory effects

Of the 80 analytes detected by LC-MS/MS, 53 displayed differential dynamics between regenerative vs degenerative VML injuries. For degenerative defects, a greater abundance of many pro-inflammatory eicosanoids such as the major lipoxygenase derived hydroxyeicosatetraenoic acids (HETEs) 5-HETE, and 12-HETE as well as cyclooxygenase (COX) derived prostaglandins including PGF_2α_, PGE_2_, PGD_2_, PGA_2_, PGI_2_ (6-keto-PGF1a) and TXB2 was detected (Figure 2B). To gain further insights into time-series variations between the two injuries, we performed non-parametric clustering of differentially detected analytes(26) (Supp. Fig. 2A). We observed variations in temporal profiles for classical eicosanoids such as prostaglandin (PGF_2α_, cluster 1) and 12-HETE (cluster 4), whereby for regenerative defects, a temporal increase in concentration was detected followed by returns to uninjured levels by 14 dpi. In contrast, PGF_2α_, and 12-HETE remained elevated in concentration for degenerative defects at 7 and 14 dpi (Figures 2C-D). In further agreement with this observation, parametric statistical analysis of analytes pooled by specific biosynthetic pathways that lead to pro-inflammatory eicosanoid production such as 5-LOX, and 12-LOX showed increases in concentration for degenerative VML injuries at longer time points driven mainly by greater and/or more prolonged local biosynthesis of PGF_2α_ (COX pathway), 5-HETE (5-LOX pathway), and 12-HETE (12-LOX pathway) (Supp. Figs. 2B-F).

To determine if the increase in eicosanoids for degenerative defects was balanced by increases in specialized pro-resolving mediators and their related pathway markers/biosynthetic intermediates (5-HEPE, 4-HDoHE, 7-HDoHE, 12-HEPE, 14-HDoHE, RvD6, PD1, 10S,17S-DiHDoHE, Mar1_n3DPA_, and LXA_4_), we plotted the ratio between classical pro-inflammatory eicosanoids (sum of TXB_2_, PGD_2_, PGE_2_, PGF_2α_, 6kPGF_1α_, and 5-, 12-, 15-HETEs) relative to detected specialized pro-resolving mediators and their related pathway markers (Figure 2E). These data revealed at 14 days, an overall stronger inflammatory response for degenerative defects when compared to regenerative defects. In contrast to the increased and sustained levels of eicosanoids, pro-resolving pathway markers/biosynthetic intermediates such as from Maresin 1 (14-HDoHE) and D-series resolvins/protectins (17-HDoHE) from clusters 2 and 5, transiently increased in abundance and returned to uninjured levels by 14 days (Figures 2F-G). The pro-resolving mediators also displayed a highly similar trajectory between degenerative or regenerative injuries (Figures 2F-G). Overall, the distinct intramuscular lipid mediator profile between injury types and time points suggests a dysregulated immune response may be driven, in part, by a relative overabundance of classical pro-inflammatory eicosanoids within degenerative VML injuries in the absence of a coordinated pro-resolving lipid mediator response. Collectively these data suggest that intervention with pro-resolving mediators to restore balance may be an effective strategy to re-establish tissue homeostasis.

### Treatment of Volumetric Muscle Loss With Maresin 1 Reduces Fibrosis and Augments Muscle Regeneration

Maresin 1 has previously been shown to reduce inflammation, inhibiting neutrophil accumulation and altering macrophage phenotype during tissue regeneration(20, 24). To evaluate whether treatment of degenerative VML injuries with exogenous Maresin 1 impacts fibrosis and muscle regeneration, we locally administered synthetic Maresin 1 (7*R*,14*S*-dihydroxydocosa-4Z,8*E*,10*E*,12*Z*,16*Z*,19*Z*-hexaenoic acid)(20) following degenerative VML injury through intramuscular injection every other day beginning at 1-dpi (Figure 3A). At 7 dpi, a significant reduction in collagen deposition was observed by Picrosirius red staining for muscles treated with Maresin 1 compared to vehicle-treated contralateral limbs (Figures 3B-C, n = 9 tissues from 9 mice, paired). Based on observed reductions in collagen deposition, we next sought to understand differences in inflammatory cell abundance. Both immunohistofluorescent stains for CD68 and flow cytometry quantifications (CD45^+^F4/80^+^) of macrophages revealed a reduction for muscles treated with Maresin 1 compared to vehicle-treated controls (Figures 3D-E, n = 7 tissues from 7 mice, paired. Supp. Figure 3A-B, n=8-10 muscles from 5 mice, unpaired). Moreover, consistent with literature showing reduced neutrophil accumulation as a result of Maresin 1 treatment(27), flow cytometry at 7 dpi revealed significant reductions in CD45^+^Ly6G^+^ neutrophils (Supp. Fig. 3A-D, n = 8-10 tissues from 5 mice, unpaired). In line with improved regeneration, we also observed a shift in the distribution of regenerating myofibers with a trend towards larger MYH3^+^ myofibers for tissues treated with Maresin 1 when compared to vehicle-treated tissues (Figure 3F, Supp. Fig. 3E, n = 7 tissues from 7 mice, paired), respectively. To determine whether the reduction in fibrosis and increase in regenerating myofiber diameters improved restoration of muscle force, we measured maximal tetanic force at 28 dpi for Maresin 1 treated tissues and vehicle-treated controls. We detected treatment with Maresin 1 yielded increases in maximal tetanic force when compared to the vehicle alone (Figure 3G-H, Supp. Fig. 3F-H, n = 7 tissues from 7 mice, paired). These results suggest that repletion of Maresin 1 reduces inflammation and fibrosis after VML and promotes a permissive microenvironment for regeneration and restoration of function.

**Figure 3.**
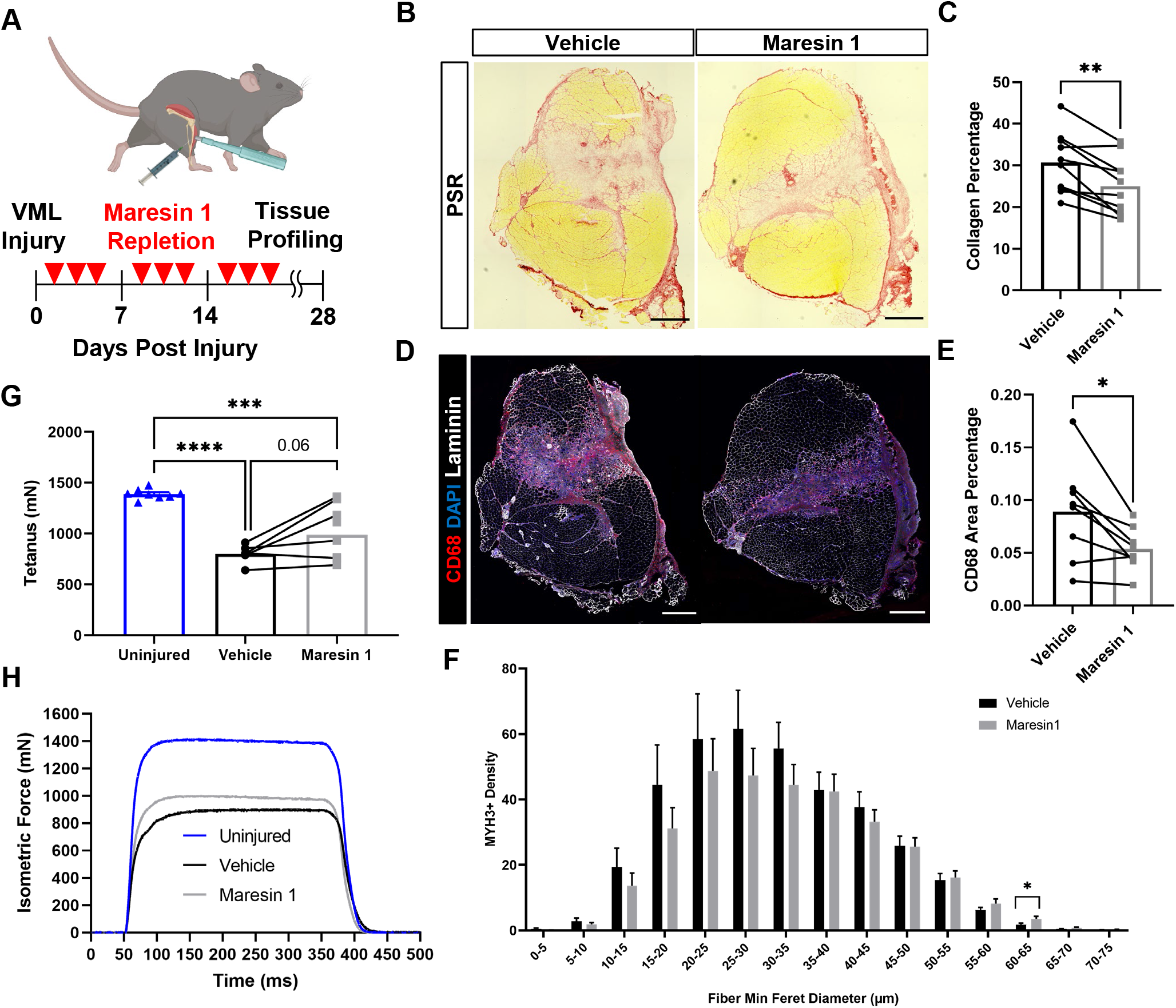
Repletion of Maresin 1 after degenerative volumetric muscle loss injury results in reductions in inflammation and fibrosis. **A**) Mice were administered a bilateral 2 mm volumetric muscle loss (VML) defect to tibialis anterior (TA) muscles and one limb received an intramuscular injection of Maresin 1 and the other limb received vehicle (saline+.01% EtOH) treatment every two days. **B)** Representative cross-sections stained with Picrosirius Red isolated 7 days after VML muscle injuries treated with vehicle or Maresin 1 treatment. n=7 mice per group, scale bar = 500 μm. **C)** Quantitation of Picrosirius Red from B) shows a reduction of collagen deposition for Maresin 1 treatment. Graphs show mean ± SEM, ** p< 0.01 by paired t-test. **D)** Representative cross-sections from muscles isolated 7 days after VML muscle injury treated with vehicle or Maresin 1 treatment. Sections are immunostained for CD68 (red), DAPI (blue) and laminin (white). n=7 mice per group, scale bar = 500 μm. **E)** Quantitation of number of macrophages (CD68^+^) from D) shows a decrease in the number of macrophages for Maresin 1 treatment. Graphs show mean ± SEM, * p< 0.05 by paired t-test. **F)** Distributions of Feret diameter for eMyHC^+^ myofibers shows increased number of smaller regenerating myofibers for vehicle treatment when compared to Maresin 1 treatment. Graphs show mean ± SEM, * p< 0.05 by unpaired t-test. n=7 mice per group **G)** Average tetanic force from muscle stimulation of uninjured (blue) tibialis anterior muscle at 28 days following 2mm volumetric muscle loss injuries treated with Saline (black) and Maresin 1 (gray) 19 days p.i. Bars show mean ± SEM and ***p<0.0005 between uninjured and VML+Maresin 1 treatment, ****p<0.0001 between uninjured and VML+Saline treatment, and p=0.0571 between VML+Maresin 1 and VML+Saline treatment by one-way ANOVA and post-hoc. n=9-6 / group. **H)** Representative force curves of uninjured tibialis anterior muscle (blue) at 28 days following 2mm injuries and Saline (black) and Maresin 1 treatment (gray).

### LGR6 Receptor Mediates Response to Maresin 1 in Muscle Stem Cells

To further probe how Maresin 1 may improve myogenesis, we sought to determine whether this lipid mediator has direct impacts on muscle stem cells (MuSCs). Previous research(28) has demonstrated that Maresin 1 selectively binds to the Leucine-rich repeat containing G protein–coupled receptor 6 (LGR6), but not other LGR receptors, such as LGR4 or LGR5. To understand whether LGR6 is expressed by MuSCs, and potential changes in expression dynamics across different MuSC states (quiescent, activated, differentiated), we assessed changes in *Lgr6* expression via RT-qPCR at 3 time points (immediately post MuSC isolation from uninjured limb muscles, following in vitro activation and culture, and 72 hours post differentiation induction using low-serum media). We used isolated MuSCs from uninjured hind limb muscles(29, 30) and observed low but detectable expression of *Lgr6* among freshly isolated MuSCs and differentiated, fused myotubes, with a nearly 65-fold increase in expression among proliferating myoblasts (Figure 4A, n = 3 wells per condition). This is consistent with previously published bulk RNA-Seq data showing upregulated *Lgr6* expression among MuSCs three days post barium chloride injury(31), and suggests that activation of the LGR6 receptor by Maresin 1 may influence proliferation(32). To test if Maresin 1 stimulated proliferation, we isolated MuSCs from uninjured limb muscles and exposed them to Maresin 1 in the presence of 5-ethynyl-2’deoxyuridine (EdU) for 24 hours. In line with our hypothesis, we observed a significant increase in EdU positive cells as a result of Maresin 1 treatment (Figure 4B, Supp. Fig. 4A, n = 4 wells per condition, unpaired).

**Figure 4.**
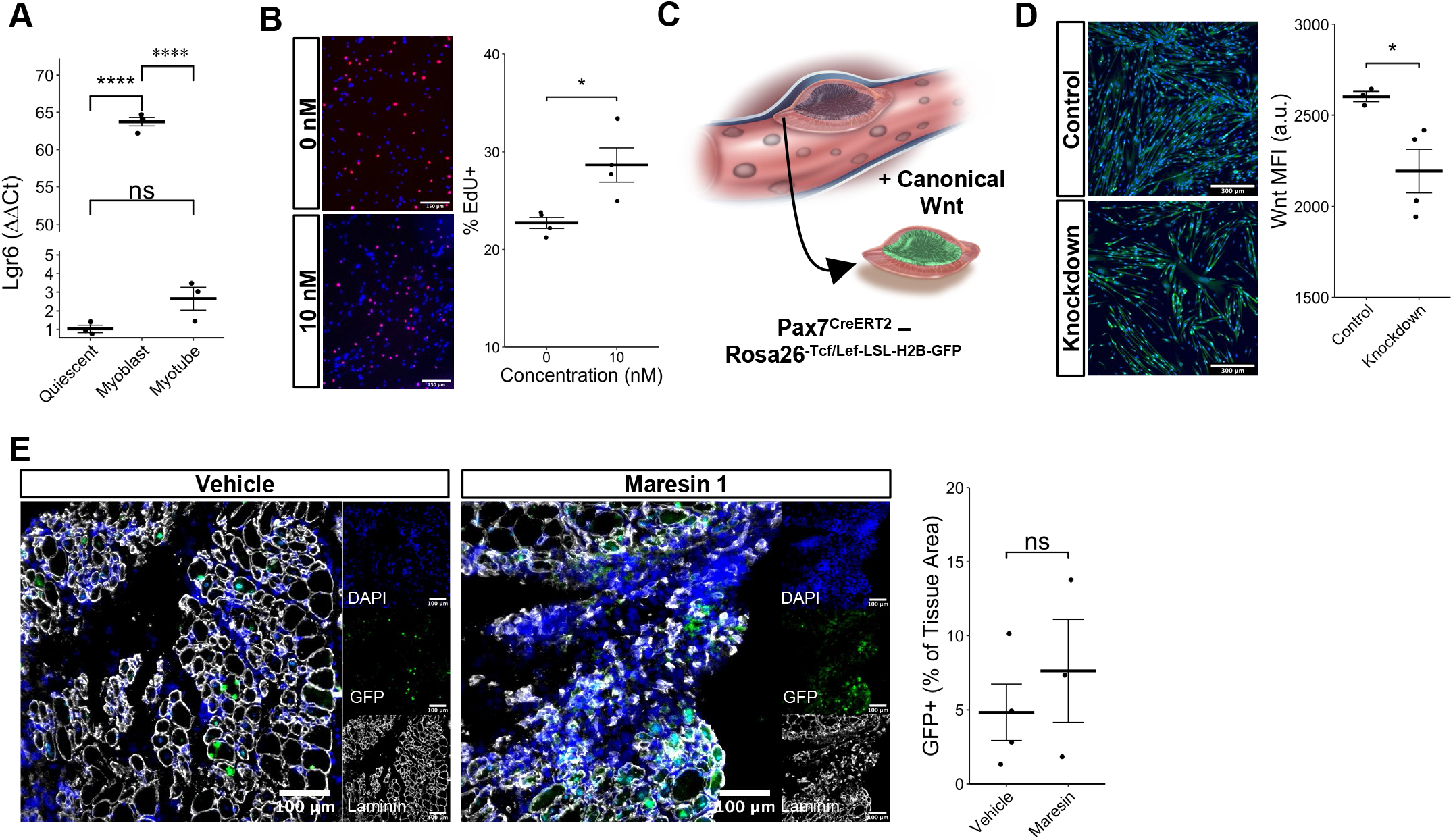
Maresin 1 impacts muscle stem cell proliferation through Lgr6 and canonical Wnt signaling. **A**) Lgr6 is highly upregulated in proliferating myoblasts by RT-qPCR. ****p<0.0001 by one-way ANOVA and BH post hoc analysis. n=3-4 wells per condition. **B)** Treatment of myoblasts with a low concentration of Maresin-1 in vitro increases proliferation based on EdU incorporation over 24 hours. *p<0.05 by one-way ANOVA with Benjamini-Hochberg post hoc analysis. n=4 wells per condition, where each well was quantified using 4 separate 10x images. Scale bars indicate 150um. **C)** Schematic of muscle stem cell lineage reporter for canonical Wnt signaling. In response to canonical Wnt, cells express green fluorescent protein in their nuclei. **D)** Wnt expression is reduced following DsiRNA knockdown of Lgr6 in primary myoblasts isolated from a P7Wnt-GFP mouse model. *p<0.05 by two-sided, two-sample t-test. n=3-4 wells per condition. Quantification results for each well were calculated using four 10x images. Scale bars indicate 300um. **E)** In vivo treatment of VML injuries with Maresin 1 slightly increases activation of canonical Wnt signaling in Pax7+ MuSCs using the P7Wnt-GFP mouse model. ns denotes p > 0.05 by two-sided, two-sample t-test. n = 3-4 tissues per condition. Scale bars indicate 100um.

Engagement of LGR6 in different cell types has been shown to stimulate several different pathways including cyclic AMP (cAMP)(32) and canonical Wnt signaling(32, 33), which have been associated with MuSC proliferation, migration, and differentiation(34, 35). To determine whether LGR6 and Maresin 1 influence canonical Wnt signaling in MuSCs, we developed a Pax7Cre^ERT2^-Rosa26^Tcf/Lef-LSL-H2B-GFP^ reporter mouse (P7Wnt) that activates a nuclear green fluorescent protein reporter in MuSCs in response to canonical Wnt signaling. We isolated MuSCs from uninjured P7Wnt hind limb muscles and used dicer-substrate short interfering RNA (Dsi-RNA) to knock down *Lrg6* (Supp. Fig. 4B). Following 72 hours in low-serum media, we observed reductions in canonical Wnt as a result of *Lgr6* knockdown via immunostaining for GFP and quantifying mean fluorescent intensity (Figure 4C, n = 4 wells per condition, unpaired). To determine whether Maresin 1 treatment increased canonical Wnt in MuSCs in vivo after degenerative VML injuries, we injured P7Wnt mice with bi-lateral degenerative VML and treated with vehicle or Maresin 1. Consistent with our in vitro results, in vivo treatment of degenerative VML injuries with Maresin 1 showed a trend towards increased GFP and canonical Wnt signaling among MuSCs at 7 dpi (Figure 4D, n = 3-4 tissues from 3 mice, unpaired). These results demonstrate that Maresin 1 influences myoblast proliferation and canonical Wnt signaling, and these effects are at least partly mediated through engagement with the LGR6 receptor.

### Single cell RNA sequencing supports increases in myogenic cells and reductions in fibrotic, lipid associated macrophages as a result of Maresin 1 repletion

To further probe the impacts of Maresin 1 treatment post VML, we performed droplet-based single cell RNA sequencing (scRNA-Seq) on viable mononucleated cells isolated from vehicle and Maresin 1 treated degenerative VML defects at 7 dpi (Figure 5A, each condition represents a pool of three tissues from three mice). We generated 2,421 and 1,421 high-quality scRNA-Seq libraries from the Maresin 1 treated and vehicle treated tissues, respectively, encompassing on average 1,708 and 1,824 genes per cell with an average read depth of 6,376 and 6,805 unique molecular identifiers (UMIs) per cell for the two treatments (Supp. Fig. 5A). SCTransform(36) was performed for normalization and variance stabilization, regressing out mitochondrial genes, followed by principal component analysis, unsupervised Louvain clustering, and Uniform-Manifold Approximation and Projection(37) to reveal 7 cell types (Figure 5A). Cluster-based cell-type annotation was performed according to the expression of known marker genes (Supp. Fig. 5B) and alignment with previously published datasets(10). Consistent with histological stains and Maresin 1-induced proliferation of myoblasts in vitro, we observed reductions in monocyte derived cells and increases in myoblasts for VML injured tissues treated with Maresin 1 (Figure 5B). Higher-resolution clustering of monocyte derived cells revealed sub-populations of dendritic cells, and two macrophage phenotypes (Figure 5C). While all three abundances decreased with Maresin 1 treatment, only the reduction in fibrotic macrophages was significant (Figure 5D). Differential gene expression across the myeloid derived cells revealed upregulated expression of genes associated with fibrotic and lipid associated macrophages(38) *(Trem2, Spp1, Apoe,* Complement) among the fibrotic macrophage subpopulation, as well as genes associated with an M2-like phenotype among the predominate other macrophage population (*Cd86, Il1b, Il1r2*) (Figure 5E). Myeloid derived TGFβ1 has a previously established role in the progression of skeletal muscle fibrosis following traumatic injuries(12), and Trem2^hi^Spp1^hi^ macrophages have reported to upregulate TGFβ1 gene expression compared to other macrophage phenotypes(39). Consistent with this, we observed slight reductions in active TGFβ1 at 7 dpi in VML-injured muscles treated with Maresin 1 compared to those treated with vehicle (Supp. Fig. 5C, n = 3-4 tissues from 3-4 mice, unpaired). Together, these results support the regenerative impact of Maresin 1 treatment being realized principally through both promoting the expansion of myoblasts and reducing immune-cell-induced fibrosis.

**Figure 5.**
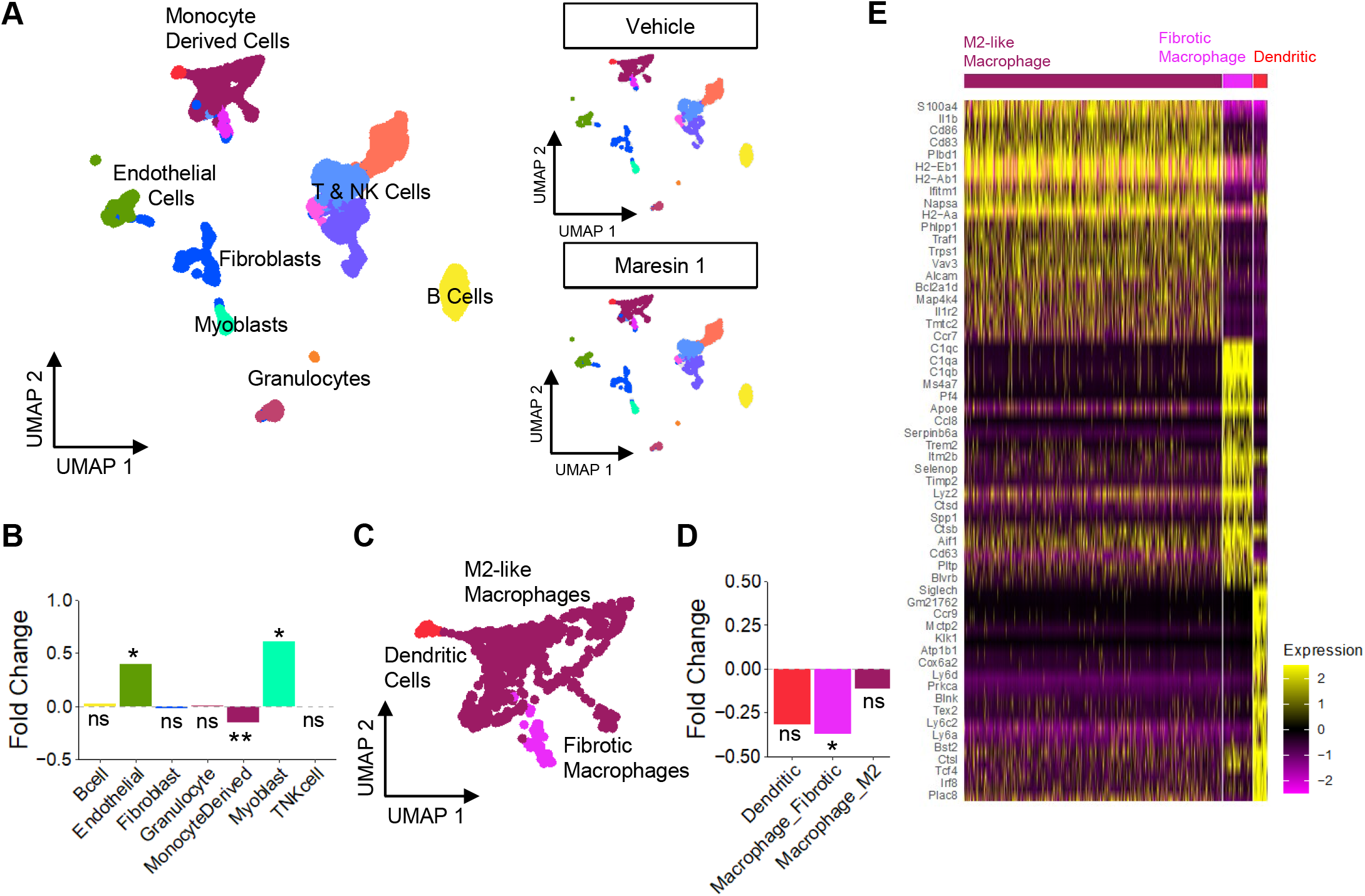
Cellular and molecular response to Maresin 1 repletion after degenerative volumetric muscle loss supports enhanced myogenic response and reduction of fibrotic macrophages. **A)** UMAP plots colored by clusters at a resolution of 0.2 and annotated into celltypes. (right) UMAP plots split by treatment. **B)** Fold changes in cell type abundance across conditions. Positive fold changes indicate increased abundance in Maresin 1 treated samples. *p<0.05, **p<0.01, ns p > 0.05 by two-sided z test for proportions. **C)** Annotation of the monocyte derived cells into two macrophage phenotypes and dendritic cells. **D)** Fold changes in abundance of monocyte derived cell types shows a significant reduction in fibrotic macrophages as a result of Maresin 1 treatment. *p<0.05 by twosided z test for proportions. **E)** Heatmap of top upregulated genes for each monocyte derived cell type.

## DISCUSSION

VML is a devastating type of acute trauma that results in fibrosis and loss of muscle function. Inadequate understanding of the drivers of these pathological outcomes has reduced the efficacy of many different types of regenerative therapies(3) and, as such, VML repair remains an unmet clinical need. Herein, we established a critical threshold model for volumetric muscle loss in murine tibialis anterior muscles. Consistent with previous observations in an analogous model in the rectus femoris(10, 25), defects below the critical threshold regenerate, while larger defects result in long-term increases in fibrosis and reductions in contractile force. Using this model of regenerative and degenerative VML, we performed metabolipidomics analysis(40) over a time course to understand signaling factors that contribute to the observed changes in fibrosis. We detected a stark imbalance of pro-inflammatory eicosanoids to pro-resolving lipid mediators in degenerative VML injuries, which correlates with our previous observations of sustained inflammation and persistent neutrophil(10) and macrophage(6) presence following degenerative VML. To determine whether restoring this balance of pro-inflammatory to pro-resolving lipids impacts regenerative outcomes, we administered the pro-resolving mediator Maresin 1 to mice following a degenerative VML injury. Administration of Maresin 1 attenuated fibrosis, reduced macrophage and neutrophil abundance, increased MuSC activation, and partially restored muscle force compared to vehicle-treated controls. Summing these results extends our understanding of muscle fibrosis and how targeting extracellular lipids can alter cell dynamics to manipulate this pathological outcome towards muscle regeneration.

The magnitude and duration of the inflammatory response after muscle injury is a critical determinant of healthy regeneration(8). After VML, the immune response becomes dysregulated(7) and contributes to fibrotic scarring. Eicosanoids are principal regulators of inflammation and we detected increases in pro-inflammatory arachidonic acid-derived eicosanoids such as LTB4, PGE_2_ and PGF_2α_ as well as other eicosanoid metabolite pathways (5-HETE, 15-HETE) for degenerative VML through all time points measured. These results contrasted with detected pro-resolving lipid mediators (RvE3, 8-oxoRvD1, LXA4, RvD6, PD1, PDX, and MaR1) that did not vary significantly with larger injury sizes. Given neutrophils and inflammatory macrophages have been demonstrated to produce pro-inflammatory mediators in injured muscle(41), and we have previously shown persistence of neutrophils in degenerative VML for weeks after injury(10), these results are consistent with increased and sustained inflammation. Our observations are distinct from muscle injuries with barium chloride(42) or cardiotoxin injection(41), where inflammation subsides quickly after injury and pro-resolving lipid mediators increase in abundance. While we observed that pro-resolving lipid mediators are expressed in VML injuries, the level at which these factors are detected did not proportionally increase with injury severity and inflammation. These results suggest that while short bursts of inflammation and transient exposure to eicosanoids such COX-derived PGE_2_ may be beneficial for muscle regeneration (43, 44), longer-term exposure to these signaling factors and concomitant immune dysregulation without balance from pro-resolving lipid mediators(45, 46) as observed in degenerative VML are detrimental to muscle regeneration.

The role of macrophage polarization towards a regenerative phenotype in guiding repair outcomes, such as the development and progression of fibrosis versus successful regeneration(47) has been well established(48). Regenerative macrophages are a significant source of pro-resolving lipid mediators after injury(41, 49), which, in combination with our results showing disproportional levels of pro-inflammatory to pro-resolving lipids following degenerative injury supports that macrophages do not effectively transition(15, 50) towards a regenerative phenotype(51) after degenerative VML(52). We speculate that the sustained inflammation and lipid imbalances post degenerative VML injury skew macrophage phenotype towards a fibrosis promoting subtype (*Spp1*^+^, *Trem2^+^, C1qc^+^, Cd63^+^*). This phenotype has also been observed in other fibrotic outcomes such as cirrhosis(53) and obesity(54). Injection of Maresin 1, a 14S-dihydroxyl-containing pro-resolving lipid mediator that is synthesized from fatty acid docosahexaenoic acid(55, 56), into degenerative VML-injured muscle reduced total and fibrotic macrophage density and collagen deposition. Since *Trem2*^+^ macrophages have recently been shown to be regulated by lipid metabolism, these results suggest that Maresin 1 treatment may restore lipid homeostasis and inhibit adoption of a macrophage pathological phenotype(57). While a deeper understanding of how macrophage phenotype is manipulated by lipid concentration and type(58), our results are in line with previous studies showing Maresin 1(59) reduced *TGF-β1*(60) and fibrotic macrophage expansion potentially through inactivation of mitogen-activated protein kinase/ERK1/2 signaling and stimulation of AMPK(61).

Maresin 1 signaling is mediated in part via engagement of the LGR6 receptor, which is expressed on numerous stem and progenitor cells, including in the skin (62), kidney (63), and mammary gland (64), and enhances proliferation, migration, and differentiation. In line with this, we observed low but detectable *Lgr6* expression in primary quiescent MuSCs, with drastic transient upregulation during activation. Additionally, we observed that Maresin 1 treatment resulted in increased numbers of MuSC derived progenitors / myoblasts after degenerative VML. These results agreed with in vivo data of MuSCs after regenerative injury(31), and further suggest Maresin 1 contributes to regenerative actions of MuSCs after injury by increasing proliferation. Given canonical Wnt signaling is a downstream target of LGR6 and induces MuSC proliferation(65), we used loss-of and gain-of-function assays to show variations in canonical Wnt activity and proliferation with changes in Maresin 1. We speculate the changes in proliferation and canonical Wnt signaling in MuSCs from Maresin 1 treatment may also be derived indirectly through differences in their ability to adhere to the matrix, given that β-catenin interacts with multiple cadherins(34, 35). Coupled together, these results show that Maresin 1 repletion can have a positive effect on ameliorating the fibrotic response as well enhance MuSC-mediated muscle regeneration after VML.

Recovery from severe muscle trauma resulting in VML is an unmet clinical need and open musculoskeletal injuries are responsible for a large fraction of hospital costs and disability payments(1)(2). The development of strategies to address lipid imbalances for this type of trauma may open new paradigms to explore coupled immuno-regenerative(43)(66) therapies.

## ACKNOWLEDGEMENTS

The authors thank the University of Michigan DNA Sequencing Core for assistance with sequencing and Jeremy Nathans for providing the *Rosa26-Tcf/Lef-LSL-H2B-GFP* strain. Research reported in this publication was partially supported by the National Institute of Arthritis and Musculoskeletal and Skin Diseases of the National Institutes of Health under Award Number P30 AR069620 (CAA and SVB), Genentech Research Award (CAA), the 3M Foundation (CAA), American Federation for Aging Research Grant for Junior Faculty (CAA), the Department of Defense and Congressionally Directed Medical Research Program W81XWH2010336 and W81XWH2110491 (CAA), a National Science Foundation CAREER award (2045977), Defense Advanced Research Projects Agency (DARPA) “BETR” award D20AC0002 (CAA) awarded by the U.S. Department of the Interior (DOI), Interior Business Center, a fellowship to J.A.C.M by the Howard Hughes Medical Institute through the James H. Gilliam Fellowships for Advanced Study program (GT15755) and the National Science Foundation Graduate Research Fellowship Program under Grant Number DGE 1256260 (J.A.L.). The content is solely the responsibility of the authors and does not necessarily represent the official views of the National Institutes of Health or National Science Foundation, the position or the policy of the Government, and no official endorsement should be inferred.

## Accession Code

GSE215808

## Author contributions

J.A.C.M, J.A.L., E.C.W., B.D.S., A.E., C.D., K.R.M., J.F.M., performed experiments. J.A.C.M, J.A.L. and B.A.Y. analyzed data. J.A.C.M., J.A.L., and C.A.A. designed the experiments. J.A.C.M., J.A.L., and C.A.A. wrote the manuscript with input from other authors.

## Materials & Methods

### Animals

C57BL/6 wild-type male and female mice (3-4 months old) were obtained from Jackson Laboratory or from a breeding colony at the University of Michigan (UM). Pax7Cre^ERT2^-Rosa26^Tcf/Lef-LSL-H2B-GFP^ mice were obtained from a breeding colony at UM and administered 5 daily 100uL intraperitoneal injections of 20mg/mL tamoxifen in corn oil. All mice were fed normal chow ad libitum and housed on a 12:12 hour light-dark cycle under UM veterinary staff supervision. All procedures were approved by the Institutional Animal Care and Use Committee (IACUC) and were in accordance with the U.S. National Institute of Health (NIH).

### Injury model

Mice were anesthetized with 5% isoflurane and maintained at 3% isoflurane. Buprenorphine analgesic was administered at 0.1 mg/kg dose via intraperitoneal injection prior to administering a volumetric muscle loss injury. The surgical area was prepared by removing hair and sterilizing through series of 70% ethanol, and betadine scrubbing. An incision of approximately 5 mm was administered to the skin to expose the tibialis anterior muscle. A full thickness volumetric muscle loss injury was administered using a sterile biopsy punch of 1 mm or 2 mm diameter to the middle of the muscle followed by closure with sutures. Animals were monitored daily for 7-10 days before removing sutures.

### Tissue sectioning

After euthanasia, uninjured or injured tibialis anterior muscles were harvested and embedded in an optical cutting temperature compound and frozen in isopentane cooled with liquid nitrogen. Cross-sections were extracted from the frozen tissue blocks using a cryotome at the midpoint of the injury and delicately placed onto positively charged glass slides.

### Picrosirius staining and quantification

Tissue sections were first fixed in 4% paraformaldehyde (PFA) for 15 minutes at room temperature. Next, the tissue sections were washed two times with 1x phosphate buffered saline (PBS) and followed by two washes with deionized (DI) water. The sections were then air dried for 20 minutes and stained with Sirius red dye for 1 hour in a humidifying chamber. Sirius red dye was washed with DI water one time for 5 minutes followed by sequential dehydration immersions in 50%, 70%, 70%, 90%, 100% ethanol solutions, and two 5 minutes incubation in xylenes at room temperature. Coverslips were mounted with Permount and whole section images were imaged using a motorized Olympus IX83 microscope. Total tissue and collagen area ratios was quantified by thresholding in FIJI or MATLAB and graphed in Graphpad.

### Metabolipidomics

C18 columns were conditioned using 15% methanol, and hexane. Elutions were performed by doing 2 washes using 100% methanol and dried using a gentle stream of nitrogen gas. After resuspending dried elutions in 50 microliters of methanol-25mM aqueous ammonium acetate (1:1), LC-MS/MS was performed in a prominence XR system (shimaduzu) using Luna C18 columns. LC-MS/MS data were analyzed using MetaboAnalyst 4.065.

### Processing of lipid abundance data

Raw lipid abundances were normalized and prepared for downstream analyses using the MetaboDiff package(67) (v0.9.5) in R (v4.2.1). Outlier samples were identified using principal component analysis (PCA) and removed, and knn imputation was repeated for the remaining samples. The data was then subjected to variance stabilization normalization (vsn) for downstream processing.

### Differential lipid abundances analysis

One-way ANOVAs (aov command in R) were performed for each lipid for the injured time points with the following design formula: Concentration ~ Condition, where Condition = {Injury + Time}, Injury = {1mm, 2mm}, and Time = {3, 7, 14 days}. Differential lipids were identified as those with p-values less than 0.05 after Benjamini-Hochberg correction.

### DPGP clustering

To cluster differentially abundant lipids by similar abundance dynamics over the time course, we used the Dirichlet process Gaussian process mixture model (DPGP v0.1)(26). Normalized imputed abundances were averaged within each condition and fold-changes were calculated between injuries (2mm over 1mm) at each time point. Fold-changes for each lipid across time points were normalized as z-scores then clustered with DPGP using default parameters with the following command:

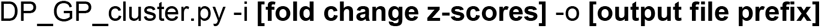

### CD68 and MYH3 Immunohistochemistry Stain and Quantification

Tissue sections were thawed and air dried in room temperature for 5 minutes followed by fixation in 100% acetone cooled to −20°C for 10 minutes or 4% paraformaldehyde in PBS at room temperature for 15 minutes. Tissues were rehydrated with 1X PBS for 5 minutes and blocked in 10% goat serum diluted in 1X PBS for 1 hour or MOM blocking reagent (Vector Labs). Primary antibodies (CD68 BioRad, MYH3 DHSB) were incubated overnight at 4°C in a humidifying chamber. Secondary antibodies (Thermo Fisher) were incubated for one hour at room temperature in a humidifying chamber. Coverslips were mounted using prolong diamond antifade. Whole section images were acquired using a Nikon A1 confocal microscope. The CD68+ area was quantified as a percentage of the full section area as using MATLAB. Myofiber regions were determined using Cellpose(68) and LabelsToROIs(69), followed by quantification of MYH3+ fibers and MYH3+ fiber measurements using MATLAB. Three sections per tissue were imaged, quantified, and averaged and graphed in Graphpad prism.

### Flow Cytometry

Mouse TAs were extracted, separately weighed using sterile surgical tools and minced using surgical scissors. Minced tissues were added to tubes containing Collagenase type II (0.2%) and Dispase II (2.5U/mL) in 10mL of DMEM, then placed on rocker in a 37°C incubator for 1 hour and mixed by pipette every 30 minutes. The enzymes were then inactivated by addition of 20% heat-inactivated fetal bovine serum (HI-FBS) in Ham’s F10 media, and the solution was passed through a 70um cell strainers, centrifuged, washed, and counted. The single cell suspension was then pelleted and re-suspended in staining buffer (PBS with 2% BSA, 2mM EDTA and 0.01% sodium azide), and plated in a 96-well round bottom plate. Cells were centrifuged at 350xg for 2.5 minutes and re-suspended in a primary antibody cocktail including CD45-APC (BioLegend), F4/80-AF488 (BioLegend), and Ly6G-APCCy7 (BioLegend) for 30 minutes on ice. Cells were then centrifuged, washed with staining buffer, then re-suspended in staining buffer containing propidium iodide for 1 minute at room temperature in the dark, centrifuged, and re-suspended in staining buffer for flow cytometry analysis. Prior to acquisition, cells were filtered through 40mm cell strainers. Single color controls were made using UltraComp eBeads (Thermo Fisher) compensation beads stained according to manufacturer’s protocol. Samples were acquired within 30 minutes on a BioRad Ze5 cytometer, and the data was processed using FlowJo (version 10) with manual compensation.

### Muscle Stem Cell Enrichment

Mouse TA muscles were extracted, and separately weighed using sterile surgical tools and placed into separate petri dishes containing ice-cold PBS. Using surgical scissors, muscle tissues were minced and Collagenase type II (0.2%) and Dispase II (2.5U/mL) were added to 10mL of DMEM per quadricep. Samples were placed on rocker in a 37°C incubator for 1.5 hours and mixed by pipette every 30 minutes. The enzymes were then inactivated by addition of 20% heat-inactivated fetal bovine serum (HI-FBS) in Ham’s F10 media. The solution was passed through a 70um cell strainers, centrifuged, and washed. Negative MuSC enrichment was performed using the Miltenyi Satellite Cell Isolation Kit for mouse according to manufacturer’s protocols or by FACS sorting for CD45-Ter119-CD31-CD11b-Sca1-B1int+CXCR4+ cells as previously described(31).

### Quantitative Real-Time PCR (qRT-PCR)

Cells were lysed directly in the plate using Buffer RLT (Qiagen) with 10uL/mL 2-mercaptoethanol following a quick PBS rinse. Cell lysates were thawed at room temperature for 30 minutes, then RNA was extracted using the Qiagen RNeasy Mini Kit according to manufacturer’s instructions. RNA purity and concentration were measured using a NanoDrop and Qubit RNA HS Assay. Within one week, cDNAs were synthesized using the SuperScript III cDNA Synthesis Kit (Thermo Fisher) according to manufacturer’s protocol. DNA quality and concentration was determined using a NanoDrop. 80-100ug cDNA template was plated in triplicate along with SYBR Green PCR MasterMix and 500nM PCR primer, then cycled 40 times starting at 95°C for 10 seconds followed by 60°C for 30 seconds on a CFX96 Real-Time thermocycler. Gene expression was quantified using the ΔΔCt method.

### In Vivo Maresin-1 Treatment

Maresin 1 (Cayman Chemicals #1268720-28-0) was aliquoted (500 ng per vial) in amber glass vials (Thermo Fisher Scientific, C4010-88AW), purged with nitrogen gas and stored at −80C. On the day of use, ethanol was evaporated using a stream of nitrogen gas and resuspended in 100 μl of sterile saline solution + 0.1% ethanol. Maresin 1 solution was protected from light and kept on ice until intramuscular administration. Mice legs were randomized to receive 100 ng of Maresin 1 (20 μl) or Vehicle (sterile saline + .1% ethanol) every two days after injury.

### In Vitro Maresin-1 Treatment

Single cell suspensions were prepared as described above in sterile conditions, followed by MACS isolation of MuSCs using the Miltenyi Satellite Cell Isolation Kit for mouse according to manufacturer’s protocols. MACS-enriched MuSCs were plated on Matrigel-coated tissue culture dishes in myoblast media (Ham’s F10 with 20% FBS, antibiotics, and fibroblast growth factor basic) and allowed to expand for up to three passages. Myoblasts were then passaged and plated in a 48-well plate with 15,000 cells seeded per well. After 24 hours, media was replaced with fresh myoblast media containing Maresin-1 (10nM) and 5-ethynyl-2’deoxyuridine (EdU) (10uM). EdU staining was performed 24 hours later using the EdU ClickIt Reaction Kit (Thermo Fisher) according to manufacturer’s instructions. Images were analyzed using MATLAB to quantify the percentage of cells positive for EdU incorporation.

### In Vitro Lgr6 Knockdown

MuSCs were MACS-enriched and cultured as described above. Following expansion in myoblast media (F10 with 10% HI FBS, bFGF, and antibiotics), cells were seeded into 12-well plates at a density of 50,000 cells per well. Lgr6 knockdown was performed using RNAiMAX (Thermo Fisher) and Lgr6 Dsi-RNA (IDT) according to manufacturer’s protocol in myoblast media without antibiotics. After 72-hours, cells were either lysed in the plate for RT-qPCR validation of knockdown efficacy, or media was replaced with myoblast media containing EdU for proliferation analysis, or with differentiation media (DMEM containing 5% horse serum and antibiotics). EdU staining was performed after 24 hours in EdU according to manufacturer’s protocols. Wnt signaling activation was assessed using cells isolated from P7Wnt mice. After 72 hours in differentiation media, cells were washed, stained with Hoechst 33342, and imaged on a Zeiss epifluorescent microscope using a 20X objective. GFP mean fluorescent intensity among GFP+DAPI+ cells was quantified using MATLAB.

### Single-cell RNA sequencing

#### Sample Preparation and Sequencing

Male and female mice received bilateral 2mm TA VML defects, which were treated with 20uL intramuscular injections of Maresin 1 (100ng in 0.1% EtOH in saline) or vehicle (0.1 % EtOH in saline) at days 1, 3, and 5 post injury. Animals were euthanized as described above at 7-dpi. TA muscles were pooled from 3 mice according to treatment, then digested into single cell suspensions as described above. Labeling with cell multiplexing oligos (CMOs) (10x Genomics) was performed according to manufacturer’s instructions (Demonstrated Protocol CG000391 Rev B, Protocol 3). Each treatment condition (vehicle and Maresin 1) was labeled with a separate CMO tag. Then equal cell numbers from each sample were pooled, stained with 7-AAD, and FACS sorted to remove dead cells and debris. Post FACS, 8,000 cells were loaded into the 10x Genomics chromium single cell controller and single cells were captured into nanoliter-scale gel bead-in-emulsions (GEMs). cDNAs were prepared using the single cell 3’ Protocol as per manufacturer’s instructions and sequenced on a NovaSeq 6000 (Illumina) with 26 bases for read1 and 98×8 bases for read2.

#### Data Processing and Analysis

10x CellRanger v7.0.0 software’s mkfastq and multi command were run with default parameters except expect-cells=8000. HD5 files were imported into R v.4.2.1 (https://www.r-project.org/) using the Seurat(70) v4.2 package and genes expressed in less than 200 cells or cells expressing less than 3 genes were removed. Seurat objects were then merged. Normalization and scaling were performed using Seurat’s SCTransform^**Error! Bookmark not defined**.^ function, regressing out mitochondrial genes. Linear dimensional reduction was performed using RunPCA, followed by FindNeighbors(dims=1:30) and RunUMAP(37) (dims=1:30). Clustering was performed using the Louvain unsupervised clustering algorithm at a resolution=0.2. Cluster marker genes were determined using Seurat’s FindAllMarkers function (only.pos = T, logfc.threshold = 1) to annotate cell types. Seurat, dittoSeq(71) and ggplot2 were used for data visualization.

### In situ functional testing

These procedures are modified from (72). Briefly, mice were anesthetized with intraperitoneal injections of tribromoethanol (250 mg/kg) and supplemental injections given to maintain an adequate level of anesthesia during the procedure. Hindlimb fur was removed with animal clippers. The TA muscle was exposed by removing the overlying skin and outer fasciae. The distal TA tendon was isolated, and the distal half of the TA was freed from adjacent muscles by carefully cutting fasciae without damaging muscle fibers. A 4–0 silk suture was tied around the distal tendon, and the tendon was severed. The animal was then placed on a temperature-controlled platform warmed to maintain body temperature at 37°C. A 25-gauge needle was driven through the knee and immobilized to prevent the knee from moving. The tendon was tied securely to the lever arm of a servomotor via the suture ends (6650LR, Cambridge Technology). A continual drip of saline warmed to 37°C was administered to the TA muscle to maintain temperature. The TA muscle was initially stimulated with 0.2 ms pulses via the peroneal nerve using platinum electrodes. Stimulation voltage and muscle length were adjusted for maximum isometric twitch force (Pt). While held at optimal muscle length (Lo), the muscle was stimulated at increasing frequencies until a maximum force (Po) was reached, typically at 200 Hz, with a one-minute rest period between each tetanic contraction. Subsequently, the same procedure was repeated, but rather than activating the muscle via the peroneal nerve, a cuff electrode was placed around the muscle for stimulation. Muscle length was measured with calipers, based on well-defined anatomical landmarks near the knee and the ankle. Optimum fiber length (Lf) was determined by multiplying Lo by the TA Lf/Lo ratio of 0.6. After the evaluation of isometric force, the TA muscle was removed from the mouse. The tendon and suture were trimmed from the muscle, and the muscle was weighed. Total muscle fiber cross-sectional area (CSA) of TA muscles was calculated by dividing muscle mass by the product of Lf and 1.06 mg/mm^3^, the density of mammalian skeletal muscle(73). Specific Po was calculated by dividing Po by CSA.

### TGFβ1 ELISA

Muscles were extracted at 7 dpi as described above and flash frozen in liquid nitrogen, then stored at −80°C. Tissues were thawed in ice-cold PBS, weighed, minced, and homogenized with 30 passes of a Dounce homogenizer in 500uL of RIPA buffer (Thermo Fisher) with a protease inhibitor cocktail (Thermo Fisher). Total protein was quantified using a Pierce BCA Assay kit (Thermo Fisher) according to manufacturer’s instructions. Active TGF**β**1 was quantified using the mouse TGF beta 1 DuoSet ELISA kit (R&D Systems) according to the manufacturer’s instructions. Absorbances were measured on a Synergy Neo microplate reader.

### Statistics

Experiments were repeated at least twice, apart from scRNA-Seq. Bar graphs show mean ± standard error unless otherwise stated. Statistical analysis was performed in GraphPad, and/or R using two-sample Student’s t-test assuming normal distribution and equal variances, one-way ANOVA, or paired-T-test, as specified in figure captions. All statistical tests performed were two-sided. Outliers were determined using the IQR method and removed from further analysis. P-values less than 0.05 were considered statistically significant.

## Competing interests

The authors declare no competing interests.

---

**Supplemental Figure 1.**
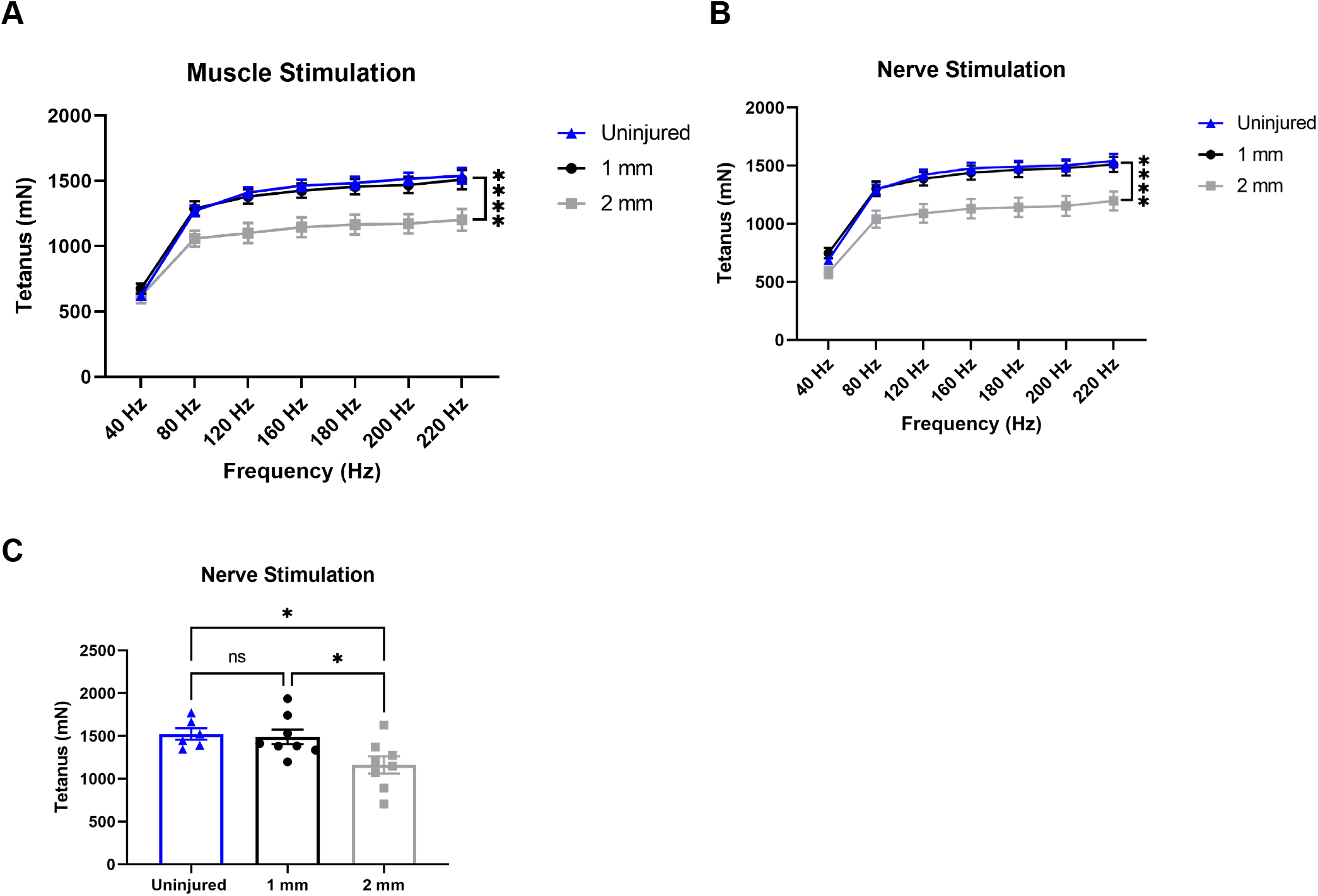
Functional assessment of response to healing or degenerative volumetric muscle loss injuries. **A-B)** Force frequency of muscle **(A)** and nerve stimulation **(B)** of uninjured, 1mm and 2 mm groups uninjured (blue) tibialis anterior muscle at 28 days following 1mm (black) or 2mm (gray) VML injuries. Points show mean ± SEM and ****p<0.0001 between uninjured and 2 mm injury groups by two-way ANOVA and post-hoc. n=6-8 / group. **C)** Average tetanic force from nerve stimulation of uninjured tibialis anterior muscle at 28 days following 1mm or 2mm VML injuries. Bars show mean ± SEM and *p<0.05 between injury types by two-way ANOVA and post-hoc. n=6-8 / group.

**Supplemental Figure 2.**
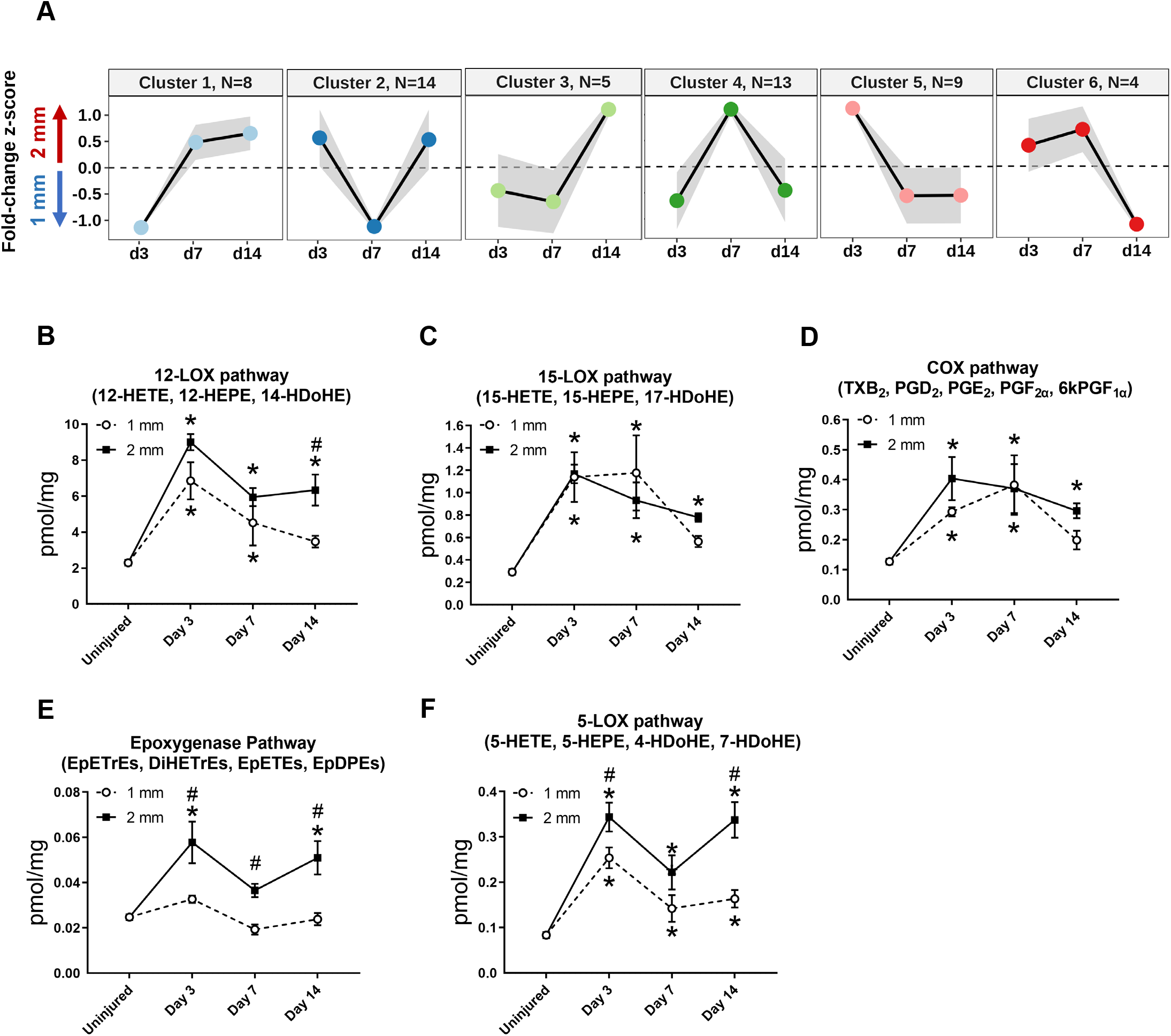
Dynamics of lipid mediators after volumetric muscle loss injury. **A)** DPGP mixture model-based clustering of mean analyte abundance fold-change z-scores across injury timepoints. Black lines are cluster means and the gray bars show 2 standard deviations around the mean. **B-F)** Changes of pooled lipid mediators metabolites per injury type and timepoint expressed in pmol/mg. Two-way paired ANOVA was used to estimate statistical differences between injury and timepoints. Graphs show mean ± SEM, * < 0.05 between timepoints of matched injury timepoint and # < 0.05 between different injury types at a certain timepoint. **B)** Sum of 12-HETE, 12-HEPE, 14-HDoHE. 12-HETE is commonly related to pro-inflammatory effects while 14-HDoHE is a known pathway marker for Maresin 1. **C)** Sum of 15-HETE, 15-HEPE, 17-HDoHE. 15-HETE is commonly related to pro-inflammatory effects while 17-HDoHE is a known pathway marker for Resolvings. **D)** Sum of PGI2 (6kPGF1a), PGF2a, PGE2, PGD2, TXB2. Prostaglandins and thromboxanes have been commonly related to pro-inflammatory effects. **E)** Sum of EpETrEs, DiHETrEs, EpETEs, EpDPEs. Cytochrome P450 epoxygenase pathway has been related to anti-inflammatory effects that remains as an understudied pathway. **F)** Sum of 5-HETE, 5-HEPE, 4-HDoHE, 7-HDoHE. 5-HETE is commonly related to pro-inflammatory effects.

**Supplemental Figure 3.**
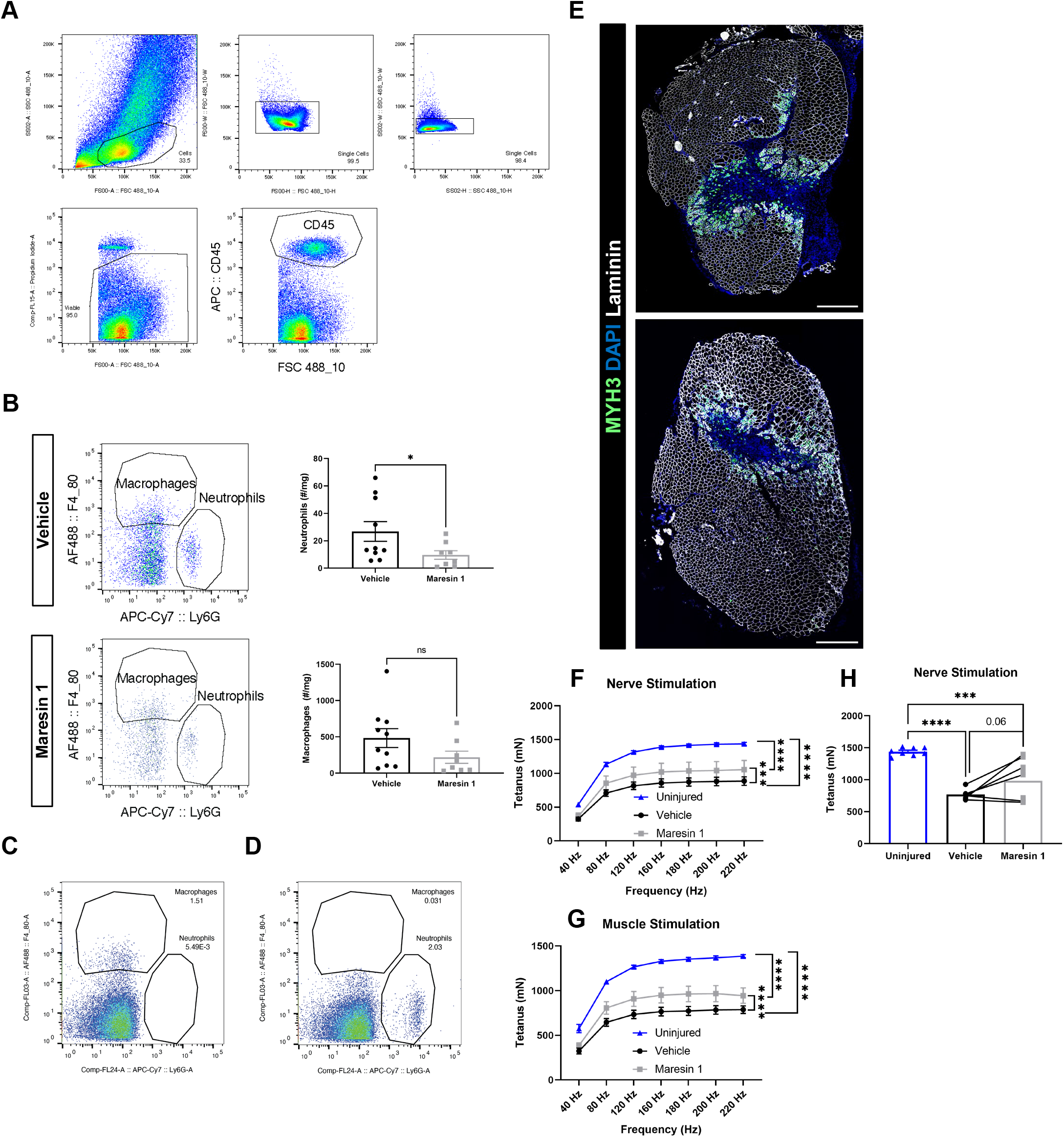
Cellular and functional profiling of response to VML injury after treatment with Maresin 1. **A)** Complete gating strategy for neutrophils and macrophages. Values indicate percentage of parent. **B)** Maresin 1 treatment significantly reduced neutrophil abundance and reduced F4/80+ macrophage abundance by flow cytometry at 7-dpi. *p<0.05 by two-sided, two-sample t-test. **C)** FMO control for Ly6G-APC-Cy7. Values indicate percentage of CD45+ cells. **D)** FMO control for F4/80-AF488. Values indicate percentage of CD45+ cells. **E)** Representative cross-sections from muscles isolated 7 days after VML muscle injury with vehicle (top) or Maresin 1 (bottom) treatment. Sections are immunostained for embryonic myosin heavy chain or eMyHC (green), DAPI (blue) and laminin (white). Scale bar = 500 μm. **F-G)** Force frequency of nerve **(F)** and muscle stimulation **(G)** of uninjured (blue) tibialis anterior muscle at 28 days following 2mm volumetric muscle loss injuries treated with Saline (black) and Maresin 1 (gray) 19 days p.i. Points show mean ± SEM and ****p<0.0001 between uninjured and vehicle, uninjured and Maresin 1 treatment, and ***p<0.001 between vehicle and Maresin 1 treatments **(F)** ****p<0.001 between vehicle and Maresin 1 treatments **(G)** by two-way ANOVA and post-hoc. n=6-9 / group. **H)** Average tetanic force from nerve stimulation of uninjured tibialis anterior muscle at 28 days following 2mm volumetric muscle loss injuries treated with Saline and Maresin 1 19 days p.i. Bars show mean ± SEM and ****p<0.0001 between uninjured and VML+Saline treatment and ***p<0.0003 between uninjured and VML+Maresin 1 treatment, and p=0.0559 between VML+Maresin 1 and VML+Saline treatment by one-way ANOVA and post-hoc. n=9-6 / group.

**Supplemental Figure 4.**
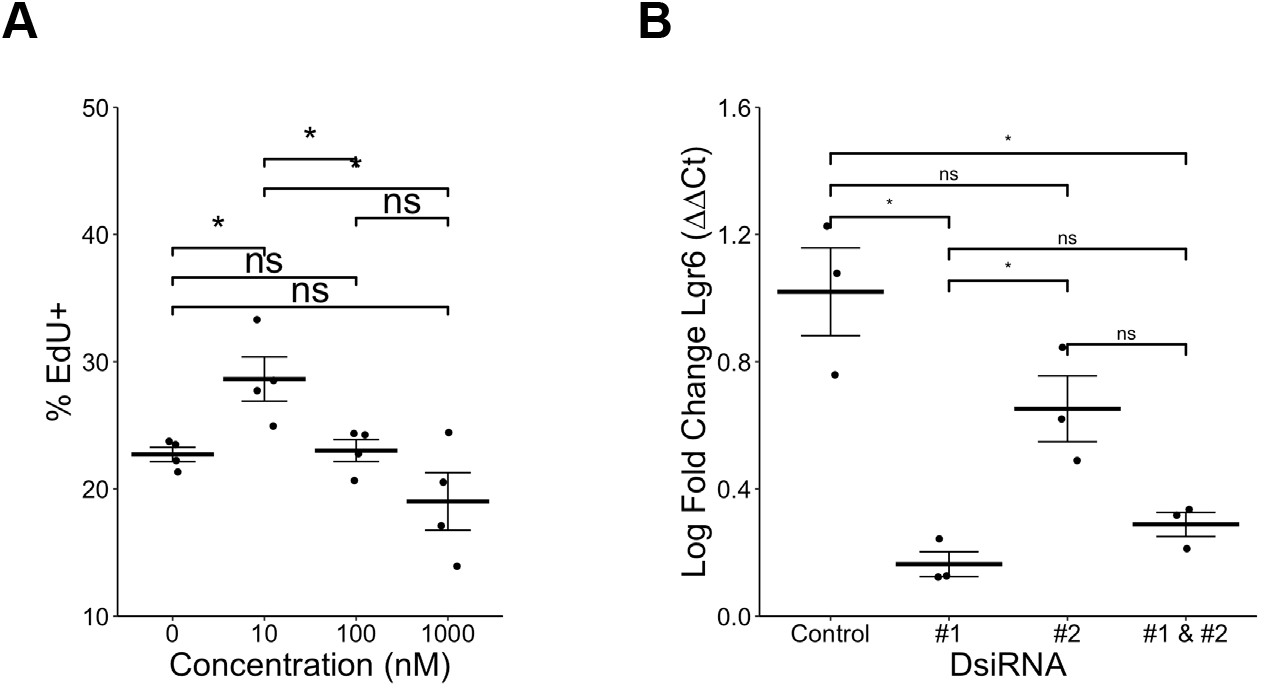
Experimental evaluation of parameters for Maresin-1 treatment of MuSCs and their progeny in vitro. **A)** Maresin-1 dose response curve showed the most impact on proliferation with treatment using 10nM. There was no statistically significant effect with higher doses. *p<0.05 by one-way ANOVA with post-hoc analysis. n=4 wells per condition, each reflecting the combined analysis of 10 images. **B)** C) Lgr6 knockdown efficiency using different DsiRNAs shows over 80% Lgr6 knockdown using construct #1. *p<0.05, ns denotes not significant by one-way ANOVA with post-hoc analysis. DsiRNA #1 was selected for further experiments.

**Supplemental Figure 5.**
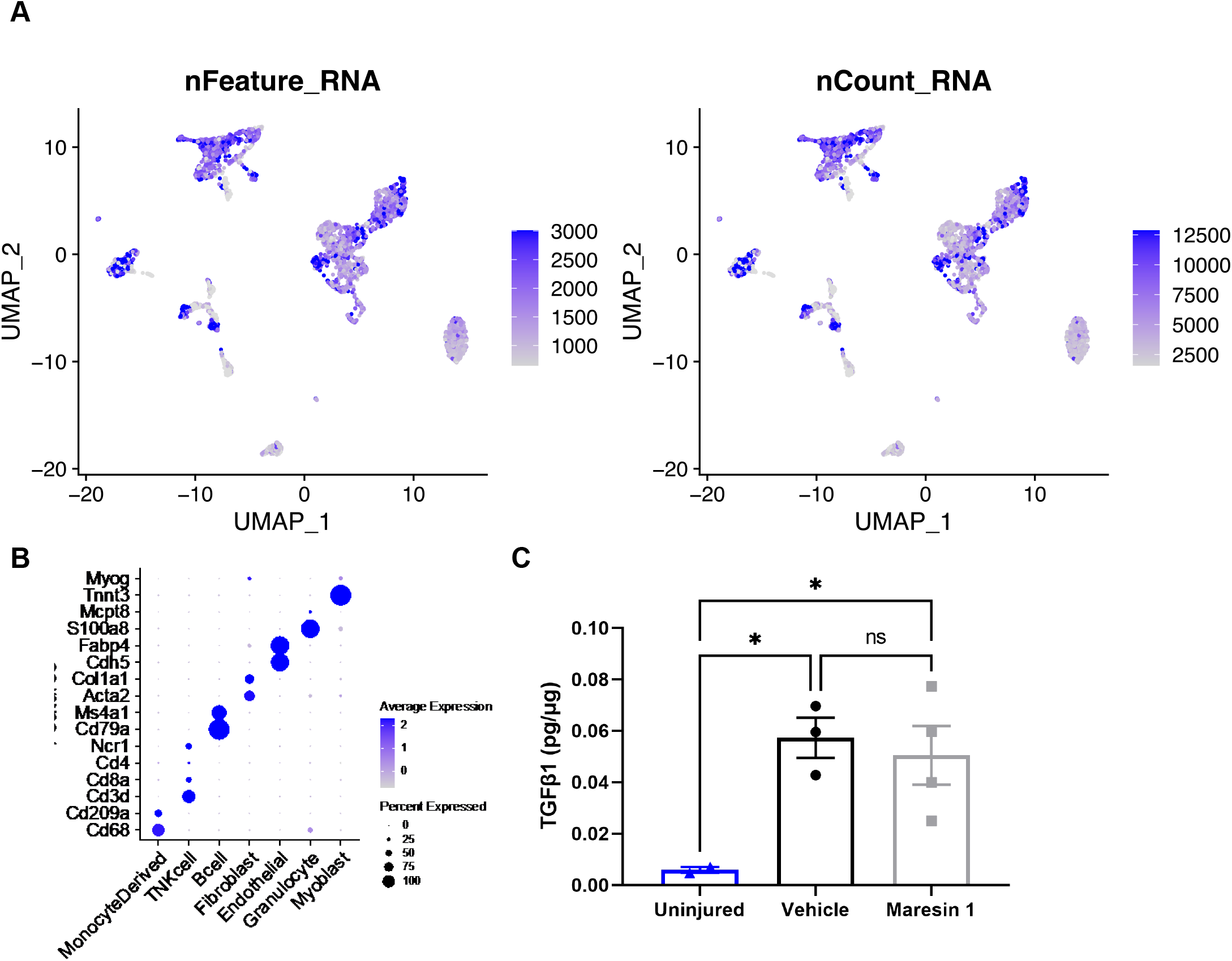
Quality control metrics for single cell sequencing after VML injury and treatment with Maresin 1. A) UMAP plots colored by the number of genes per cell (left) and the number of unique molecular identifiers per cell (right). B) Dot plot of known marker gene expression for each cell type. C) ELISA assay targeting TGFβ1 in VML injured TAs muscles. Maresin 1 and vehicle treatments were administered every 2 days throughout a 7 days time course after injury. Points show mean ± SEM and *p<0.05 between uninjured and vehicle, uninjured and Maresin 1 treatments by one-way ANOVA and post-hoc. n=3-4 / group.

## Notes

### Competing Interest Statement

The authors have declared no competing interest.

