## Supplemental Table 1 for "Maresin 1 Repletion Improves Muscle Regeneration After Volumetric Muscle Loss"

**Supplementary Table 1: Metabolite concentration (pmol/g) for different pathways in uninjured, 1 mm VML, and 2 mm (TA) tissue homogenates.**

| | | | 3 days | | 7 days | | 14 days | |
|---|---|---|---|---|---|---|---|---|
| Pathway | Analytes | Uninjured | 1 mm | 2 mm | 1 mm | 2 mm | 1 mm | 2 mm |
| Cyclooxygenase | PGE1 | ND | 2.491967 ± 0.36384 | 3.563911 ± 0.433393 | 2.832362 ± 1.313814 | 1.971901 ± 0.773613 | 0.953444 ± 0.503735 | ND |
| | PGF1a | 0.535076 ± 0.180779 | 0.938721 ± 0.118269 | 0.839611 ± 0.292882 | 1.085034 ± 0.126875 | 1.064194 ± 0.076653 | 0.732907 ± 0.085164 | 1.151028 ± 0.086089 |
| | 15-keto PGE1 | ND | ND | ND | ND | ND | ND | ND |
| | 13,14dhPGE1 | ND | ND | ND | ND | ND | ND | ND |
| | 13,14dh-15k-PGE1 | ND | 0.214298 ± 0.088746 | 0.432757 ± 0.158348 | 0.497531 ± 0.083907 | 0.378132 ± 0.130879 | 0.209127 ± 0.089098 | ND |
| | D17-PGE1 | ND | ND | ND | ND | ND | ND | ND |
| | Bicyclo PGE1 | ND | ND | 0.489826 ± 0.167917 | ND | ND | ND | ND |
| | 6-keto PGE1 | ND | ND | ND | ND | ND | ND | ND |
| | 2,3-dinor PGE1 | ND | ND | ND | ND | ND | ND | ND |
| | 19(R)-hydroxy PGE1 | ND | ND | ND | ND | ND | ND | ND |
| | 15(R)-PGE1 | ND | ND | 1.652695 ± 0.265267 | ND | ND | ND | ND |
| | TXB2 | 14.53873 ± 2.919347 | 21.95302 ± 1.606664 | 23.55804 ± 8.679987 | 27.84298 ± 4.964594 | 19.97881 ± 1.624087 | 13.38531 ± 1.819587 | 20.08715 ± 1.970618 |
| | 12(S)-HHTrE | 82.25863 ± 6.316511 | 217.0924 ± 14.61969 | 350.7983 ± 23.9393 | 291.2639 ± 48.26767 | 338.1022 ± 16.16505 | 189.9027 ± 18.03918 | 334.1237 ± 41.55752 |
| | PGD2 | 15.87406 ± 1.521426 | 40.27702 ± 3.932453 | 50.09019 ± 7.244723 | 88.52967 ± 20.13218 | 77.35252 ± 16.32767 | 39.10384 ± 7.797462 | 64.02356 ± 10.12217 |
| | PGE2 | 62.94715 ± 5.959717 | 162.7386 ± 14.01119 | 257.6128 ± 53.57187 | 213.092 ± 40.91186 | 193.7438 ± 37.77063 | 100.0821 ± 14.07895 | 148.1741 ± 12.32078 |
| | PGF2a | 17.74357 ± 0.914807 | 26.46533 ± 2.653353 | 31.6344 ± 3.341197 | 36.26276 ± 3.866642 | 36.00637 ± 1.644134 | 24.66226 ± 1.752778 | 37.52463 ± 2.648139 |
| | 6kPGF1a | 16.20208 ± 3.042057 | 26.41505 ± 2.522163 | 41.13438 ± 4.376048 | 47.21256 ± 8.374545 | 41.96629 ± 3.308935 | 16.95567 ± 3.672448 | 26.55724 ± 3.108848 |
| | PGJ2 | 2.173939 ± 0.453536 | 4.287008 ± 0.407516 | 6.639899 ± 0.793945 | 9.442145 ± 1.784219 | 7.790942 ± 1.182364 | 4.708802 ± 0.862526 | 9.297291 ± 1.396738 |
| | PGA2 | 2.585839 ± 0.573045 | 5.961919 ± 0.480145 | 10.1377 ± 1.382585 | 12.27537 ± 2.251229 | 9.793398 ± 1.620396 | 6.146677 ± 1.406903 | 11.94994 ± 1.937559 |
| | 15-keto PGE2 | 4.207772 ± 0.413594 | 4.064619 ± 0.503655 | 4.755405 ± 0.786642 | 7.560881 ± 1.031173 | 7.364869 ± 1.376132 | 6.115615 ± 0.680623 | 8.447479 ± 0.534354 |
| | 15-keto PGF2a | 4.875812 ± 0.310888 | 9.891208 ± 0.683537 | 13.97421 ± 2.376933 | 12.83475 ± 2.296754 | 10.96629 ± 2.158315 | 7.493059 ± 0.93359 | 9.905688 ± 1.027914 |
| | 13,14dh-15k-PGE2 | 3.264495 ± 0.233447 | 12.94356 ± 1.264529 | 16.18909 ± 1.667532 | 11.25377 ± 2.052485 | 10.95112 ± 1.934415 | 5.307699 ± 0.706315 | 6.79748 ± 0.698089 |
| | 13,14dh-15k-PGD2 | 5.730804 ± 0.769661 | 7.186827 ± 0.592957 | 10.05923 ± 0.800203 | 11.48803 ± 2.415374 | 8.274531 ± 1.335679 | 4.506501 ± 0.98848 | 7.600597 ± 0.676319 |
| | 19(R)-OH PGF2a & 20-OH PGF2a | ND | ND | ND | ND | ND | ND | ND |
| | 8-isoPGF2a & 11bPGF2a | 0.629944 ± 0.213506 | 1.365891 ± 0.157586 | 1.886652 ± 0.258854 | 1.331845 ± 0.531493 | 2.33648 ± 0.094882 | 1.539167 ± 0.253293 | 1.505948 ± 0.50259 |
| | 11dh-2,3-dinor TXB2 | 0.322661 ± 0.117694 | 0.433705 ± 0.11355 | 0.179732 ± 0.060456 | ND ± ND | ND ± ND | ND ± ND | ND ± ND |
| | D12-PGJ2 | 1.089603 ± 0.195831 | 2.89577 ± 0.29872 | 3.259005 ± 0.428461 | 5.508135 ± 1.09742 | 4.270727 ± 0.744761 | 2.096904 ± 0.385363 | 4.268572 ± 0.646119 |
| | tetranor PGEM | ND | 0.132611 ± 0.059732 | ND | ND | ND | ND | ND |
| | 15d-D12,14-PGJ2 | ND | ND | ND | ND | ND | ND | ND |
| | Bicyclo PGE2 | ND | 0.953373 ± 0.436989 | ND | 0.624513 ± 0.223769 | ND | ND | ND |
| | 19(R)-OH PGE2 & 20-OH PGE2 | ND | ND | ND | ND | ND | ND | ND |
| | 2,3-dinor TXB2 | ND | 0.685567 ± 0.299423 | 1.778565 ± 0.288181 | ND | 1.079272 ± 0.115007 | 0.952389 ± 0.389003 | 1.441668 ± 0.493783 |
| | 11dh-TXB2 | ND | ND | ND | ND | ND | ND | ND |
| | 13,14dh-15k-PGF2a | ND | ND | ND | ND | ND | ND | ND |
| | 6,15-diketo PGFa | ND | ND | ND | ND | ND | ND | ND |
| | iPF-VI | ND | ND | ND | ND | ND | ND | ND |
| | TXB3 | ND | 0.475633 ± 0.195142 | 1.339813 ± 0.494586 | ND ± ND | ND ± ND | ND ± ND | ND ± ND |
| | PGE3 | 2.305874 ± 0.590913 | 2.502209 ± 0.248714 | 3.743294 ± 0.385793 | 3.302097 ± 0.549565 | 2.781189 ± 0.62726 | 1.538134 ± 0.285391 | 1.878501 ± 0.05774 |
| | PGF3a | ND | ND | ND | 0.38678 ± 0.020334 | ND | ND | ND |
| | PGD3 | ND | ND | ND | ND | ND | ND | ND |
| | 11dh TXB3 | ND | ND | ND | ND | ND | ND | ND |
| | 15d-D12,14-PGJ3 | ND | ND | ND | ND | ND | ND | ND |
| | sum | 237.2861 ± 24.72676 | 552.3663 ± 45.65857 | 835.7496 ± 112.1951 | 784.6271 ± 142.2959 | 776.173 ± 89.01086 | 426.3923 ± 54.72988 | 694.7346 ± 79.78537 |
| | Δ change | 0 ± 25.01288 | 315.0803 ± 44.9667 | 598.4635 ± 111.2673 | 547.3411 ± 142.3096 | 538.887 ± 89.05483 | 189.1063 ± 22.0008 | 457.4485 ± 80.77761 |
| 5-LOX | 9(S)-HOTrE | 28.13713 ± 2.401094 | 49.04578 ± 4.668874 | 58.29138 ± 8.027431 | 36.25603 ± 4.883079 | 41.66373 ± 4.153014 | 54.63865 ± 3.273427 | 73.37781 ± 3.121311 |
| | 9-OxoOTrE | 15.29822 ± 1.532393 | 19.73577 ± 0.978258 | 21.21616 ± 0.898883 | 13.58437 ± 2.593341 | 17.02492 ± 1.905426 | 24.2398 ± 1.991497 | 32.96426 ± 2.29982 |
| | 5(S)-HETrE | 0.641192 ± 0.122483 | 1.651296 ± 0.222119 | 2.00838 ± 0.261049 | 1.548028 ± 0.280345 | 1.976539 ± 0.40163 | 1.428627 ± 0.194471 | 3.364226 ± 0.378728 |
| | 5-HETE | 34.44025 ± 4.517922 | 137.9783 ± 8.711948 | 199.1843 ± 16.39475 | 99.91466 ± 20.073 | 133.8505 ± 16.51268 | 84.18443 ± 8.843736 | 190.6682 ± 27.19441 |
| | 5-oxoETE | 11.39126 ± 1.20267 | 25.89372 ± 2.493538 | 41.77412 ± 7.39832 | 19.35099 ± 4.523874 | 33.52741 ± 3.410227 | 26.2783 ± 3.304714 | 58.23399 ± 13.77704 |
| | LTB4 | 1.245387 ± 0.560459 | 1.428768 ± 0.586858 | 4.638772 ± 0.53314 | ND | 4.983473 ± 0.577223 | 1.739935 ± 0.758308 | 3.8035 ± 1.837697 |
| | 12-OxoLTB4 | ND | ND | ND | ND | ND | ND | ND |
| | 5-HEPE | 11.45911 ± 1.413296 | 23.56888 ± 2.39594 | 32.20938 ± 4.990271 | 17.72839 ± 3.151027 | 18.61485 ± 0.962803 | 16.05231 ± 1.306198 | 22.35398 ± 1.27134 |
| | LTB5 | ND | ND | ND | 0.238811 ± 0.089038 | ND | 0.431602 ± 0.127718 | 0.643312 ± 0.236831 |
| | 4-HDoHE | 35.20168 ± 2.975049 | 85.08591 ± 7.802655 | 99.68481 ± 13.01842 | 42.07771 ± 7.952423 | 66.58801 ± 9.006675 | 61.87164 ± 4.766302 | 114.9655 ± 10.99656 |
| | 7-HDoHE | 2.538678 ± 0.660292 | 9.252362 ± 1.018548 | 12.68562 ± 1.715405 | 4.452458 ± 0.956431 | 7.217722 ± 0.852462 | 5.124983 ± 0.949135 | 9.49006 ± 1.677714 |
| | sum | 140.3529 ± 15.38566 | 353.6408 ± 28.87874 | 471.6929 ± 53.23767 | 235.1515 ± 44.50256 | 325.4472 ± 37.78214 | 275.9903 ± 25.51551 | 509.8649 ± 62.79145 |
| | Δ change | 0 ± 20.27101 | 213.2879 ± 28.87874 | 331.34 ± 294.1328 | 94.79854 ± 160.2752 | 185.0942 ± 53.93872 | 135.6374 ± 25.38779 | 369.512 ± 128.1007 |
| 8-LOX | 8-HETE | 25.3469 ± 6.861517 | 93.14095 ± 19.26615 | 103.1384 ± 4.904674 | 77.10856 ± 18.86373 | 71.63837 ± 8.698124 | 45.53639 ± 4.955301 | 73.58396 ± 4.245809 |
| | 8-HEPE | 5.710808 ± 0.479515 | 12.0606 ± 2.033192 | 19.08686 ± 1.971355 | 11.32699 ± 2.160788 | 13.84576 ± 1.930551 | 11.5536 ± 1.794465 | 13.31785 ± 2.010408 |
| | sum | 31.0577 ± 7.341032 | 105.2015 ± 21.29934 | 122.2253 ± 6.876029 | 88.43555 ± 21.02452 | 85.48413 ± 10.62868 | 57.08999 ± 6.749766 | 86.90182 ± 6.256217 |
| | Δ change | 0 ± 7.341032 | 74.14385 ± 21.29934 | 91.16758 ± 6.876029 | 57.37785 ± 21.02452 | 54.42643 ± 10.62868 | 26.03228 ± 2.710771 | 55.84411 ± 6.256217 |
| 12-LOX | 9-HODE | 196.0035 ± 25.94695 | 450.0682 ± 44.57243 | 532.5914 ± 76.75486 | 431.006 ± 41.44746 | 443.3317 ± 15.94707 | 482.1621 ± 35.43303 | 777.5551 ± 119.569 |
| | 9-OxoODE | 318.8974 ± 24.56657 | 566.1251 ± 70.19947 | 570.8342 ± 89.67895 | 441.9057 ± 46.72869 | 380.9001 ± 66.56678 | 613.5087 ± 103.8389 | 612.107 ± 70.16632 |
| | 8(S)-HETrE | 4.735886 ± 0.611556 | 15.73981 ± 1.700484 | 18.86834 ± 1.401807 | 6.771765 ± 1.043452 | 12.15568 ± 1.422275 | 11.40948 ± 1.82738 | 21.42498 ± 1.894086 |
| | 11(R)-HEDE | 5.957872 ± 0.646236 | 24.58353 ± 2.769678 | 36.56846 ± 4.040438 | 15.07159 ± 3.960952 | 20.01263 ± 2.627438 | 13.51103 ± 1.947299 | 22.82707 ± 2.699604 |
| | 12-HETE | 1519.465 ± 426.0061 | 4916.774 ± 506.854 | 6373.862 ± 156.1738 | 3755.518 ± 852.7616 | 4507.317 ± 303.4065 | 2702.394 ± 383.9268 | 4617.513 ± 730.6691 |
| | 5(S),12(S)-DiHETE | 5.407379 ± 0.702952 | 15.86703 ± 0.842529 | 24.10695 ± 2.221273 | 16.58708 ± 3.926715 | 23.73092 ± 2.817835 | 13.42724 ± 1.343418 | 26.12311 ± 5.398971 |
| | tetranor 12-HETE | 4.798079 ± 1.742342 | 11.23169 ± 0.974645 | 13.85788 ± 1.466823 | 17.38687 ± 4.671815 | 9.915233 ± 0.949648 | 7.731704 ± 0.216901 | 7.512382 ± 0.878238 |
| | 12-OxoETE | ND | ND | 77.40581 ± 2.476746 | ND | 73.85217 ± 26.88201 | 34.94635 ± 14.73031 | 75.97865 ± 34.16499 |
| | 12-HEPE | 554.8689 ± 90.34904 | 1055.348 ± 121.4143 | 1563.343 ± 290.4054 | 679.1964 ± 165.9676 | 1056.391 ± 92.36137 | 904.9185 ± 134.3101 | 1250.562 ± 139.1507 |
| | 14-HDoHE | 222.9699 ± 69.40596 | 1004.863 ± 233.5216 | 1066.338 ± 61.21159 | 853.6872 ± 172.0145 | 695.9837 ± 81.85112 | 327.8657 ± 32.86824 | 470.3079 ± 21.70322 |
| | sum | 2833.104 ± 639.9777 | 8060.6 ± 982.8492 | 10277.78 ± 685.8317 | 6217.13 ± 1292.523 | 7223.59 ± 594.8321 | 5111.875 ± 710.4423 | 7881.911 ± 1126.294 |
| | Δ change | 0 ± 646.7432 | 5227.496 ± 1002.078 | 7444.673 ± 685.8317 | 3384.027 ± 1311.577 | 4390.487 ± 594.8321 | 2278.771 ± 285.3205 | 5048.808 ± 1126.294 |
| 15-LOX | 13-HODE | 1871.093 ± 359.1395 | 3771.053 ± 457.0939 | 4257.306 ± 198.6977 | 3936.705 ± 735.1959 | 3292.798 ± 283.2126 | 3391.65 ± 130.2083 | 4710.167 ± 180.0066 |
| | 13-OxoODE | 431.6364 ± 52.73821 | 627.4252 ± 23.11416 | 696.9474 ± 81.89387 | 559.1848 ± 51.60369 | 582.8574 ± 66.09269 | 785.8199 ± 46.15931 | 1285.41 ± 155.8027 |
| | 13(S)-HOTrE | 129.274 ± 45.76725 | 204.2811 ± 47.84847 | 196.6014 ± 24.07971 | 379.7862 ± 86.78558 | 203.4648 ± 27.88182 | 192.0698 ± 9.110597 | 225.8504 ± 34.46885 |
| | 13(S)-HOTrE(g) | ND | ND | ND | ND | ND | ND | ND |
| | 15(S)-HEDE | 4.250451 ± 1.324336 | 24.23076 ± 6.074965 | 25.21326 ± 1.555287 | 16.56356 ± 4.051463 | 14.19759 ± 2.509535 | 9.305937 ± 1.129274 | 13.20535 ± 0.729961 |
| | 15-OxoEDE | ND | 2.81673 ± 0.809395 | 3.767324 ± 1.242503 | ND | 1.940448 ± 0.690861 | ND | 4.73368 ± 1.260942 |
| | 15-HETE | 148.8894 ± 37.41444 | 579.5814 ± 81.44297 | 662.2214 ± 25.73291 | 619.8295 ± 125.6084 | 573.5082 ± 75.72338 | 318.9659 ± 21.0416 | 533.9221 ± 30.42944 |
| | 15-OxoETE | 9.346658 ± 2.465824 | 14.64923 ± 1.862624 | 6.489069 ± 2.416603 | ND ± ND | 5.99542 ± 2.166914 | ND ± ND | 9.619351 ± 1.919678 |
| | 8(S),15(S)-DiHETE | ND | 0.667283 ± 0.304886 | ND | 1.111253 ± 0.257655 | 0.596191 ± 0.20147 | ND | ND |
| | 5(S),15(S)-DiHETE | 0.159892 ± 0.092591 | 0.281886 ± 0.139917 | ND | 0.390513 ± 0.10262 | ND | 0.158598 ± 0.048485 | ND |
| | 15(S)-HEPE | 107.3012 ± 28.71886 | 318.3741 ± 78.89831 | 321.8667 ± 41.82554 | 490.5413 ± 113.1574 | 296.1292 ± 45.13428 | 181.217 ± 13.58752 | 173.4386 ± 21.90966 |
| | 5(S),15(S)-DiHEPE | ND | ND | 0.74857 ± 0.330973 | 0.654264 ± 0.148247 | 0.577123 ± 0.20299 | ND | 0.623351 ± 0.229797 |
| | 17-HDoHE | 34.57842 ± 11.88337 | 146.4062 ± 36.36658 | 178.669 ± 15.1883 | 144.8517 ± 28.00688 | 115.9242 ± 10.82668 | 52.01221 ± 5.221559 | 69.78053 ± 7.056268 |
| | sum | 2736.529 ± 539.5443 | 5689.767 ± 733.9562 | 6349.83 ± 392.9634 | 6149.618 ± 1144.918 | 5087.988 ± 514.6432 | 4931.199 ± 226.5066 | 7026.751 ± 433.8139 |
| | Δ change | 0 ± 539.9622 | 2953.238 ± 734.1121 | 3613.301 ± 393.3519 | 3413.089 ± 1147.617 | 2351.459 ± 514.7295 | 2194.67 ± 91.80481 | 4290.221 ± 434.1871 |
| Epoxygenase | 9(10)-EpOME | 39.17392 ± 3.549138 | 69.50146 ± 1.419018 | 70.47135 ± 24.46948 | 36.31697 ± 13.18198 | 52.52802 ± 18.21599 | 49.97423 ± 12.98732 | ND |
| | 12(13)-EpOME | 19.44482 ± 2.079992 | 32.63155 ± 0.980555 | 46.53996 ± 5.351743 | 29.94902 ± 4.575822 | 37.36401 ± 2.952332 | 34.0012 ± 2.902771 | 41.91121 ± 3.295647 |
| | 9,10-DiHOME | 18.67946 ± 3.472983 | 14.45801 ± 0.785647 | 22.13135 ± 2.327373 | 24.48568 ± 2.261555 | 22.68399 ± 3.212073 | 21.89038 ± 1.950122 | 33.22617 ± 5.669846 |
| | 12,13-DiHOME | 18.35166 ± 2.876344 | 16.42241 ± 1.171458 | 23.48214 ± 2.403102 | 22.67261 ± 1.880628 | 25.843 ± 2.781899 | 22.52642 ± 2.451236 | 41.58788 ± 11.21316 |
| | 5(6)-EpETrE | 2.001098 ± 0.253832 | 4.727364 ± 0.494843 | 6.824515 ± 1.097338 | 2.868822 ± 0.82486 | 4.060311 ± 0.35494 | 2.914907 ± 0.332516 | 5.921104 ± 0.829734 |
| | 8(9)-EpETrE | ND | ND | 1.065149 ± 0.361776 | ND | ND | 0.692891 ± 0.322915 | 1.443109 ± 0.555539 |
| | 11(12)-EpETrE | 3.085592 ± 0.33191 | 7.333158 ± 0.404991 | 12.011 ± 2.118878 | 4.875388 ± 1.223129 | 7.697901 ± 0.866598 | 5.384579 ± 0.616952 | 10.84936 ± 1.321166 |
| | 14(15)-EpETrE | 5.96335 ± 0.960042 | 4.93719 ± 1.542152 | 8.215703 ± 2.135155 | 1.815546 ± 0.819672 | 2.305804 ± 0.842614 | 1.333475 ± 0.645963 | 2.313217 ± 0.920659 |
| | 5,6-DiHETrE | ND | ND | ND | ND | ND | ND | ND |
| | 8,9-DiHETrE | ND | 0.155505 ± 0.069341 | 0.46634 ± 0.032446 | 0.253668 ± 0.04093 | 0.205298 ± 0.072177 | 0.161577 ± 0.07 | 0.551316 ± 0.281191 |
| | 11,12-DiHETrE | 1.672206 ± 0.228308 | 2.124363 ± 0.225765 | 3.540041 ± 0.701233 | 2.390185 ± 0.387172 | 3.072976 ± 0.48756 | 2.032122 ± 0.197182 | 4.824692 ± 1.255412 |
| | 14,15-DiHETrE | 3.91321 ± 0.887106 | 4.23466 ± 0.237727 | 7.018333 ± 1.016305 | 4.191561 ± 0.353586 | 6.099917 ± 0.755122 | 3.857449 ± 0.404749 | 8.688321 ± 2.437834 |
| | 20-HETE | ND | ND | ND | ND | 15.74908 ± 5.460279 | ND | ND |
| | 8(9)-EpETE | ND | ND | ND | ND | ND | ND | ND |
| | 11(12)-EpETE | ND | ND | ND | ND | ND | ND | ND |
| | 14(15)-EpETE | ND | ND | ND | ND | ND | ND | 1.863218 ± 0.629178 |
| | 17(18)-EpETE | ND | 0.717791 ± 0.293284 | 1.564991 ± 0.107729 | 0.695445 ± 0.241803 | 1.185165 ± 0.0957 | 0.770284 ± 0.199699 | 1.215961 ± 0.085332 |
| | 18-HEPE | 16.21113 ± 0.884296 | 25.36115 ± 3.207142 | 28.857 ± 1.602703 | 30.87253 ± 3.653449 | 32.91731 ± 1.452933 | 26.20247 ± 2.466958 | 29.28525 ± 3.600808 |
| | 7(8)-EpDPE | ND | 0.646777 ± 0.17863 | 1.009153 ± 0.078484 | 0.321628 ± 0.118323 | 0.707095 ± 0.077908 | 0.703658 ± 0.096907 | 1.118431 ± 0.141898 |
| | 10(11)-EpDPE | 1.825434 ± 0.324787 | 1.986819 ± 0.211246 | 3.60684 ± 1.178744 | 1.651115 ± 0.260029 | 1.745196 ± 0.339495 | 1.630596 ± 0.662944 | 2.466374 ± 0.354572 |
| | 13(14)-EpDPE | 0.513419 ± 0.172987 | 1.158365 ± 0.507922 | 2.924846 ± 1.119064 | ND | 1.86269 ± 0.242579 | 0.908475 ± 0.373193 | 2.755927 ± 0.411785 |
| | 16(17)-EpDPE | ND | 0.802225 ± 0.336533 | ND | ND | 1.022783 ± 0.143595 | 0.564829 ± 0.245877 | 1.227495 ± 0.434913 |
| | 19(20)-EpDPE | 2.272919 ± 0.295959 | 6.617484 ± 0.891209 | 8.970206 ± 1.273834 | 4.116209 ± 0.712767 | 5.104096 ± 0.39798 | 3.202505 ± 0.317346 | 5.840651 ± 0.452418 |
| | sum | 133.1082 ± 16.31768 | 193.8163 ± 12.95746 | 248.6989 ± 47.37538 | 167.4764 ± 30.5357 | 222.1546 ± 38.75178 | 178.752 ± 27.24464 | 197.0897 ± 33.8911 |
| | Δ change | 0 ± 16.77979 | 60.70806 ± 13.24302 | 115.5907 ± 47.74275 | 34.36815 ± 31.84644 | 89.04641 ± 33.86393 | 45.64382 ± 10.94171 | 63.98147 ± 57.99221 |
| Specialized pro-resolving mediators | LXA4 | 0.092623 ± 0.035484 | 0.411891 ± 0.039896 | 0.453387 ± 0.13382 | 0.13004 ± 0.063677 | 0.503203 ± 0.181223 | 0.151949 ± 0.065652 | 0.769801 ± 0.213638 |
| | LXA5 | ND | ND | ND | ND | ND | ND | ND |
| | LXB4 | ND | ND | ND | ND | ND | ND | ND |
| | 15-epi LXA4 | ND | ND | ND | ND | ND | ND | ND |
| | 15-oxo LXA4 | ND | ND | ND | ND | ND | ND | ND |
| | RvE1 | ND | ND | ND | ND | ND | ND | ND |
| | RvE2 | ND | ND | ND | ND | ND | ND | ND |
| | RvE3 | ND | ND | ND | ND | ND | ND | ND |
| | RvD5(n-3DPA) (7,17-DiHDoPE) | ND | ND | ND | ND | ND | ND | ND |
| | PD1(n-3, DPA) | ND | ND | ND | ND | ND | ND | ND |
| | MaR1(n-3DPA) | 1.342639 ± 0.510614 | ND | ND | ND | ND | 1.248728 ± 0.36879 | ND |
| | RvD1 & AT-RvD1 | 2.506282 ± 1.055978 | ND | ND | ND | ND | 1.716217 ± 0.266733 | 2.981461 ± 1.393127 |
| | 8-oxoRvD1 | ND | ND | ND | ND | ND | ND | ND |
| | 17-oxoRvD1 | ND | ND | ND | ND | ND | ND | ND |
| | RvD2 | ND | ND | ND | ND | ND | ND | ND |
| | RvD3 | ND | ND | ND | ND | ND | ND | ND |
| | AT-RvD3 | 1.245896 ± 0.295023 | 1.91096 ± 0.512534 | 1.529411 ± 0.205818 | ND | 0.996605 ± 0.399229 | ND | ND |
| | RvD4 | ND | ND | ND | ND | ND | ND | ND |
| | RvD5 | ND | ND | ND | ND | ND | ND | ND |
| | RvD6 (4,17-DiHDoHE) | 0.423202 ± 0.13362 | 0.677681 ± 0.066564 | 1.397186 ± 0.368778 | 0.761841 ± 0.146753 | 0.920146 ± 0.101592 | 0.566283 ± 0.035309 | 1.687143 ± 0.411976 |
| | PD1 | 8.835957 ± 3.280358 | 18.41183 ± 5.181173 | 15.64948 ± 1.313472 | 17.18365 ± 2.954572 | 9.759962 ± 2.456608 | 4.724716 ± 0.508888 | 7.456593 ± 1.314655 |
| | PDX (10S,17S-DiHDoHE) | 14.90069 ± 5.120719 | 31.98153 ± 8.322141 | 29.94415 ± 2.836109 | 35.09376 ± 5.496087 | 19.56019 ± 4.187239 | 8.993092 ± 0.728518 | 14.15731 ± 2.476874 |
| | AT-PD1 | ND | ND | ND | ND | ND | ND | ND |
| | 22-OH-PD1 | ND | ND | ND | ND | ND | ND | ND |
| | Maresin1 | ND | ND | ND | ND | ND | ND | ND |
| | 7(S)-Maresin1 | ND | ND | ND | ND | ND | ND | ND |
| | sum | 29.34729 ± 10.4318 | 53.39389 ± 14.12231 | 48.97361 ± 4.857997 | 53.16929 ± 8.661089 | 31.7401 ± 7.32589 | 17.40099 ± 1.973889 | 27.05231 ± 5.810271 |
| | Δ change | 0 ± 10.4318 | 24.0466 ± 14.97522 | 19.62632 ± 6.657511 | 23.822 ± 9.823918 | 2.392814 ± 8.240354 | -11.9463 ± 0.883615 | -2.29498 ± 7.591695 |
| Non-enzymatic | 9-HETE | ND | ND | ND | ND | ND | ND | ND |
| | 11-HETE | 153.8461 ± 17.35617 | 425.7819 ± 15.09435 | 653.9455 ± 31.42412 | 424.2041 ± 95.36895 | 459.4422 ± 28.67588 | 251.213 ± 25.59287 | 472.3782 ± 63.14737 |
| | 9-HEPE | 20.08613 ± 3.605392 | 41.61214 ± 7.199263 | 64.02293 ± 10.45603 | 23.29644 ± 6.318992 | 37.81449 ± 5.395287 | 36.31026 ± 7.286782 | 53.19582 ± 8.80214 |
| | 11-HEPE | 19.86415 ± 2.529652 | 35.7747 ± 3.29768 | 46.69397 ± 4.850812 | 35.61862 ± 6.721848 | 36.22517 ± 3.530335 | 26.01929 ± 3.378014 | 31.7966 ± 2.179641 |
| | 8-HDoHE | 8.9899 ± 1.299762 | 22.57201 ± 2.191793 | 28.67599 ± 6.170418 | 13.05164 ± 1.94399 | 18.69917 ± 0.251392 | 18.37315 ± 2.184397 | 33.60916 ± 4.966495 |
| | 10-HDoHE | 21.3954 ± 3.70608 | 64.62225 ± 9.443455 | 76.91772 ± 6.765119 | 42.85821 ± 7.70072 | 45.47846 ± 4.316635 | 30.26566 ± 2.561505 | 49.61808 ± 6.012652 |
| | 11-HDoHE | 8.845363 ± 1.866856 | ND | ND | ND | ND | 16.13314 ± 2.271076 | 28.04014 ± 2.246671 |
| | 13-HDoHE | 28.37805 ± 3.419724 | 71.92171 ± 6.361867 | 105.7202 ± 9.572847 | 58.45436 ± 9.683334 | 61.64736 ± 6.228373 | 36.97689 ± 3.471541 | 68.86174 ± 9.23755 |
| | 16-HDoHE | 28.48859 ± 2.17375 | 52.19897 ± 5.738472 | 69.55665 ± 9.158807 | 21.6113 ± 9.842381 | 40.31782 ± 3.123035 | 40.56246 ± 3.839883 | 76.20299 ± 8.739308 |
| | 20-HDoHE | 37.351 ± 3.039047 | 95.0934 ± 8.51291 | 122.0956 ± 18.04219 | 57.50005 ± 5.7698 | 81.24025 ± 6.626336 | 65.06483 ± 5.525898 | 133.152 ± 15.06105 |
| | sum | 327.2447 ± 38.99643 | 809.5771 ± 57.83979 | 1167.629 ± 96.44034 | 676.5947 ± 143.35 | 780.8649 ± 58.14727 | 520.9187 ± 56.11197 | 946.8548 ± 120.3929 |
| | Δ change | 0 ± 38.99643 | 482.3324 ± 65.7999 | 840.3838 ± 105.9315 | 349.35 ± 156.0857 | 453.6202 ± 58.14727 | 193.674 ± 22.53511 | 619.6101 ± 120.3929 |

**Values are mean ± SEM of 5 mice/group. ND = Below limits of detection of the assay in ≥ 50% of samples.**
